# Mechanism-guided engineering of a minimal biological particle for genome editing

**DOI:** 10.1101/2024.07.23.604809

**Authors:** Wayne Ngo, Julia T. Peukes, Alisha Baldwin, Zhiwei Wayne Xue, Sidney Hwang, Robert R. Stickels, Zhi Lin, Ansuman T. Satpathy, James A. Wells, Randy Schekman, Eva Nogales, Jennifer A. Doudna

## Abstract

The widespread application of genome editing to treat or even cure disease requires the delivery of genome editors into the nucleus of target cells. Enveloped Delivery Vehicles (EDVs) are engineered virally-derived particles capable of packaging and delivering CRISPR-Cas9 ribonucleoproteins (RNPs). However, the presence of lentiviral genome encapsulation and replication components in EDVs has obscured the underlying delivery mechanism and precluded particle optimization. Here we show that Cas9 RNP nuclear delivery is independent of the native lentiviral capsid structure. Instead, EDV-mediated genome editing activity corresponds directly to the number of nuclear localization sequences on the Cas9 enzyme. EDV structural analysis using cryo-electron tomography and small molecule inhibitors guided the removal of ∼80% of viral residues, creating a minimal EDV (miniEDV) that retains full RNP delivery capability. MiniEDVs are 25% smaller yet package equivalent amounts of Cas9 RNPs relative to the original EDVs, and demonstrated increased editing in cell lines and therapeutically-relevant primary human T cells. These results show that virally-derived particles can be streamlined to create efficacious genome editing delivery vehicles that could simplify production and manufacturing.

**SIGNIFICANCE STATEMENT:** Our results highlight the importance of understanding how virally-derived particles function to eliminate unnecessary viral proteins and create more efficacious and easier-to-produce delivery vehicles for therapeutic genome editing.

## Introduction

CRISPR-Cas9-mediated genome editing has enabled genetic therapies including an approved treatment for sickle cell disease.^1^ To advance the utility of genome editing in larger patient populations, however, efficient methods are needed to deliver editing enzymes into diseased cells in the body. Enveloped delivery vehicles (EDVs) are virally-derived particles that can package and transport genome editing ribonucleoproteins (RNPs) into cells in culture and *in vivo*. These particles are programmable when engineered to display both a fusogen and antibody fragments on their surface.^2,3^ While attractive as a delivery strategy for CRISPR-Cas9 RNPs, the structure and delivery mechanism of EDVs have yet to be determined.

Derived from HIV-1 lentiviral vectors, EDVs could employ multiple mechanisms of protein and nucleic acid nuclear delivery. Lentiviruses package nucleic acids and associated proteins, including nucleocapsid, into a proteinaceous capsid core structure that assembles during virion maturation.^4–7^ After HIV-1 virions escape from endosomes in infected cells, the capsid cores protect the RNA genome and associated proteins from innate immune detection, traveling along microtubules to deposit their contents into the nucleus by translocation across the nuclear pore.^8–10^ Some HIV-1 components, including the matrix protein, contain nuclear localization signals (NLSs) that bind to host proteins, such as importin α, for transport through the nuclear pore complex.^11^ Other HIV-1 proteins, such as the integrase, may use both the capsid and NLSs for nuclear delivery.^12^ Because the Cas9 RNPs packaged in EDVs comprise both nucleic acids and NLS-containing proteins, their mechanism of EDV-mediated nuclear delivery has been unclear.

Here we determined the components that are necessary for EDV-mediated genome editing. We discovered that although the capsid structure assembles in a subset of EDV particles, it does not transport Cas9 RNPs into the nucleus. Instead, NLS peptides engineered into the Cas9 protein confer nuclear entry and can be tuned to improve delivery efficiency. Furthermore, mechanism-guided engineering enabled simplification of the EDV design, creating miniEDV particles with only 22% of the original viral residues while achieving up to 2.5-fold higher editing potency compared to the original EDVs in primary human T cells. Understanding the functional components of virally-derived particles paves the way towards more effective and readily manufacturable genome editing therapies.

## Results

### The EDV capsid core does not mediate nuclear delivery of Cas9 RNPs

We showed previously that inhibiting the capsid core with a pre-clinical small molecule, GS-CA1, did not reduce EDV editing activity.^2^ This preliminary result suggested that the capsid core is not essential for nuclear transport of Cas9 RNPs, a surprising finding given the central role of the capsid structure in lentivirus cargo nuclear localization. To explore this further, we tested two additional small molecule inhibitors of the capsid core, lenacapavir and PF-3450074 (PF74) (Fig. 1a).^13–15^ We produced EDVs packaging Cas9 RNPs that target a prematurely truncated luciferase reporter gene (C205ATC).^16^ HIV-1 lentiviral vectors packaging a transgene encoding Cas9 enzymes and the same guide RNA were used as a positive control. The particles were incubated with HEK-293T cells expressing the truncated luciferase reporter in either the presence or absence of the inhibitors. Editing leads to insertions and deletions that can restore the luciferase reporter reading frame. We found that luciferase expression was specific to cleavage at the luciferase locus, proportional to the dose of EDVs and detectable 48 h after transduction (*SI Appendix,* Fig. S1). In the presence of increasing concentrations of lenacapavir, a clinically-approved HIV-1 inhibitor that impairs cargo delivery by stabilizing the core (Fig. 1b),^13,14^ no decrease in EDV-mediated induction of reporter cell luminescence occurred (Fig. 1c). Similarly, incubation of cells with increasing concentrations of the capsid core destabilizer PF74 (Fig. 1d)^15^ also had no effect on EDV-mediated luminescence (Fig. 1e). Parallel experiments with lentiviral vectors encoding analogous components (Cas9 and sgRNA against the luciferase reporter gene) showed dose-dependent loss of reporter cell luminescence, consistent with inhibitor prevention of lentivirus-mediated nuclear delivery (Fig. 1f, g). Together these results support the conclusion that the capsid core is not needed for Cas9 RNP delivery by EDVs.

**Figure 1.**
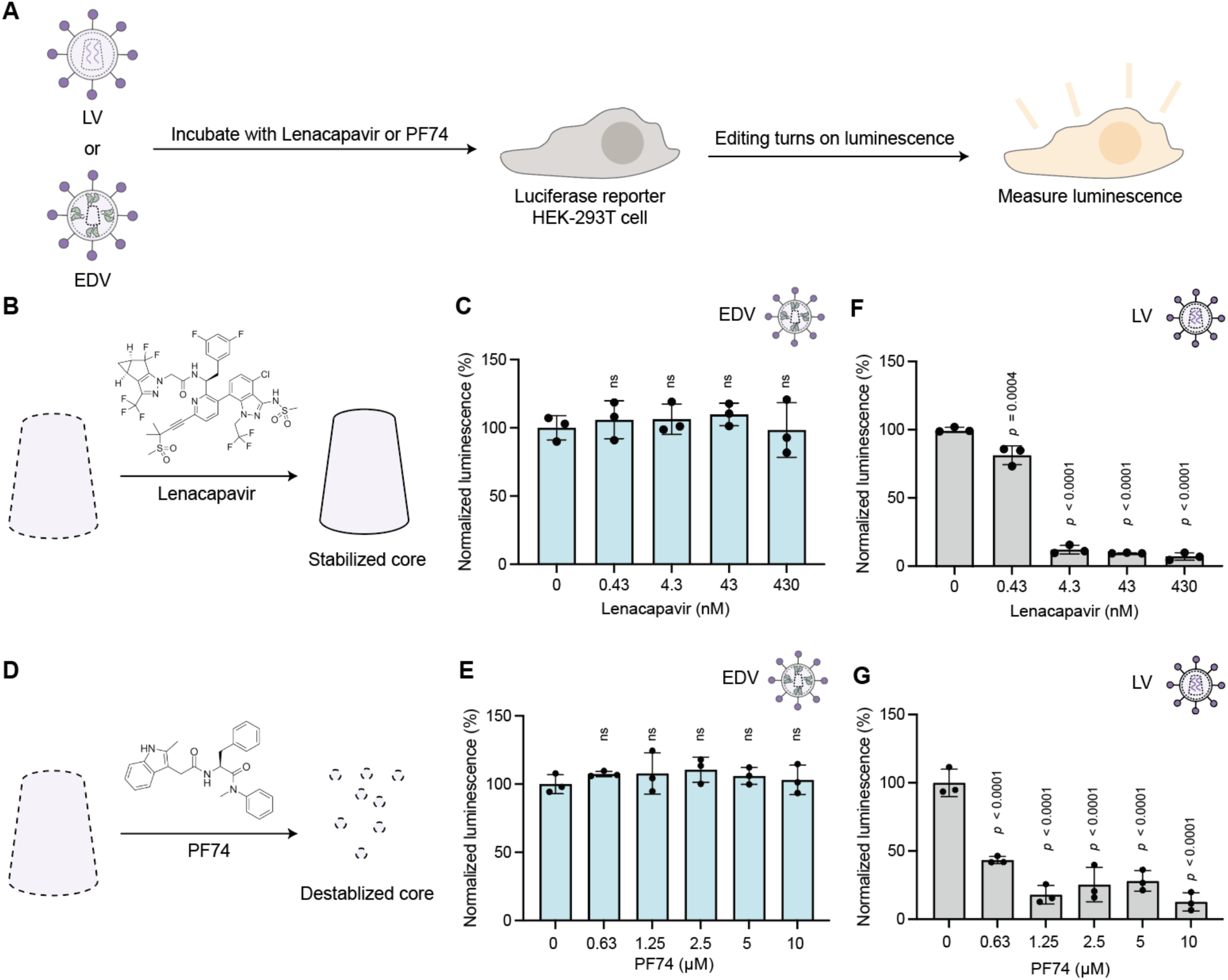
Small molecule inhibitors that disrupt the capsid core do not impact EDV editing. (a) Schematic of small molecule inhibition experiments. EDVs or lentivirus (LV) were incubated with luciferase reporter HEK-293T cells in the presence of Lenacapavir or PF74. The luminescence of the reporter cells were recorded after incubation. (b) Schematic showing that Lenacapavir stabilizes the capsid core. (c) Lenacapavir did not inhibit EDVs compared to the DMSO control (0 nM). (d) Schematic showing that PF74 destabilizes the capsid core. (e) PF74 did not inhibit EDVs compared to the DMSO control (0 nM). (f) Lenacapavir inhibited LVs compared to the DMSO vehicle control (0 nM). (g) PF74 inhibited LVs compared to the DMSO vehicle control (0 nM). Data were normalized to the DMSO control. *P*-values were calculated using an ordinary one-way ANOVA with Dunnett’s multiple comparisons test. Mean ± standard deviation of n = 3 biological replicates. Significant *p*-values as indicated. Non significance indicated by “n.s.”

As the luminescence produced by the reporter cells depends on both nuclear entry and Cas9 editing, we directly tested whether Cas9 nuclear entry required the capsid core. We incubated HEK-293T cells with EDVs and PF74 for 24 h, isolated cell nuclei and used Western blots to determine the relative amounts of Cas9 enzymes or capsid associated with the nucleus (*SI Appendix,* Fig. S2a). PF74 was used in this experiment because lenacapavir has been shown to stall capsid cores on the cytosolic side of nuclear pores, leading to their co-isolation with the nuclear fraction.^17^ We confirmed successful nuclear isolation by monitoring nuclear localization of EZH2 and cytosolic localization of GAPDH (*SI Appendix,* Fig. S2b). The 24 kDa mature capsid protein decreased in the nuclear fraction in the presence of 10 µM PF74, while the amount of Cas9 enzyme remained consistent across all PF74 concentrations (*SI Appendix,* Fig. S2b). Both the Gag-Cas9 polyprotein (220 kDa) and Cas9 (160 kDa) were present in the nuclear fractions. The presence of Gag-Cas9 in the nuclear fractions is surprising because it was assumed that editing enzymes needed to be liberated from viral structural proteins by viral protease cleavage to enable nuclear entry.^2,18–20^ Our results suggested that the liberation of Cas9 enzymes by protease cleavage may not be necessary for nuclear association. These results show that Cas9 RNP delivery into the nucleus is independent of the EDV capsid core.

### EDVs form capsid cores that do not encapsulate Cas9

We wondered why Cas9 RNP nuclear entry was independent of the capsid core. We began by testing whether EDVs contain mature capsid cores as observed in lentivirus. After purification by iodixanol cushion ultracentrifugation to remove contaminating proteins, EDVs and lentiviral vectors were visualized in their native hydrated states using cryogenic electron tomography.^21–24^ Three-dimensional tomograms of the EDVs and lentiviral vectors (Supplementary movies 1 and 2) revealed spherical particles with a lipid bilayer in each case (Fig. 2a). Surface glycoproteins appeared as dark spots densely coating the lipid bilayer exterior. We quantified and compared the proportion of mature particles (with a capsid core), immature particles (concentric rings of proteins under the lipid bilayer) and unknown particles. Consistent with previous reports, both EDVs and lentiviral particles were of similar size (∼125 nm diameter) and contained multiple morphologies of the mature capsid core (*SI Appendix,* Fig. S3a-d).^25^ A fraction of EDVs (29%) and lentiviral vectors (51%) were mature with a capsid core (Fig. 2b). Immature EDVs (36%) and lentiviral vectors (18%) were also visible (Fig. 2b). The remaining particles (∼30%) could not be categorized and were presumed to be other types of vesicles or broken particles (Fig. 2b). The higher proportion of immature EDVs compared to lentiviral vectors was confirmed in Western blots. Western blots showed that in lentiviral vectors harvested at 30, 48 or 72 h after transfection, the mature 24 kDa capsid protein was more abundant than the uncleaved (55 kDa) Gag polyprotein (Fig. 2c). In contrast, a similar analysis of EDVs showed the 55 kDa Gag polyprotein to be more abundant than the 24 kDa capsid species at all time points (Fig. 2c).

**Figure 2.**
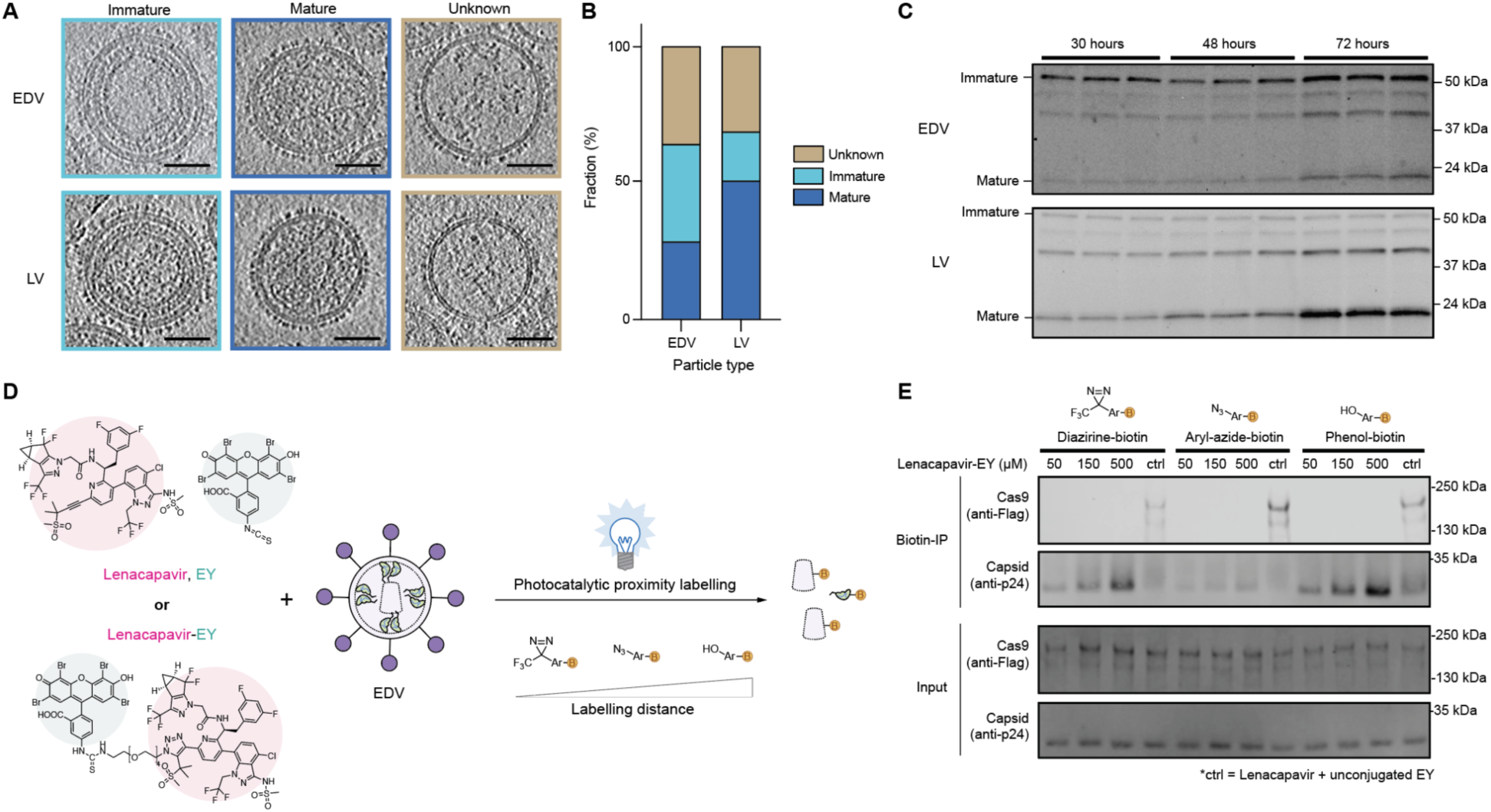
EDVs produce capsid cores that do not encapsulate Cas9. (a) Representative two-dimensional slices in the XY plane from cryogenic-electron tomograms of EDVs and LVs that are immature, mature, or unknown. Scale bars are 50 nm. (b) Fraction of EDVs (n = 498) and LVs (n = 374) that are mature, immature, or unknown. (c) Western blot showing the fraction of immature capsid protein (55 kDa) compared to mature capsid protein (24 kDa) in EDVs and LVs harvested 30, 48 or 72 h after transfection (as indicated). Three independent batches of EDVs and LVs were harvested (one per lane). Partially mature forms of the capsid protein are also visible in the blot. (d) Schematic of photocatalytic proximity labeling experiment. The Lenacapavir-eosin Y (EY) conjugate or unconjugated EY and Lenacapavir were incubated with EDVs and allowed to bind. Different photo-probes were then added to enable biotinylation of proximal proteins upon blue light illumination. Biotinylated proteins were isolated using biotin enrichment (biotin-IP). (e) Western blot demonstrating the amount of biotinylated Cas9 or mature capsid upon lenacapavir-EY-mediated photocatalytic proximity labeling. The proximity ligation experiments were repeated twice with similar results.

This observation led us to examine whether Gag protein maturation differences resulted from structural differences between immature EDVs and lentiviral vectors. We used subtomogram averaging and alignment to compare the immature capsid domains of EDVs and lentiviral vectors to those of published HIV-1 structures. This analysis revealed that the immature capsid domains of both EDVs and lentiviral vectors matched the HIV-1 structure (PDB: 5L93)^22^ with root mean square deviations of 1.2 Å and 1.7 Å respectively (*SI Appendix,* Fig. S3e-f). Together, these data show that immature capsid domains in EDVs were structurally indistinguishable from lentiviral vectors.

We next tested whether EDV editing activity is independent of the capsid core because it does not encapsulate Cas9. To test this, we used photocatalytic proximity labeling with a lenacapavir-eosin Y conjugate to map proteins located near the capsid core components (Fig. 2d). Examination of the published co-structures of lenacapavir and capsid (PDB: 7RJ4) suggested that the alkyne group on lenacapavir is not involved in binding and could potentially be used for small molecule conjugation.^26^ We incubated the lenacapavir-eosin Y or unconjugated lenacapavir and eosin Y with EDVs, then added one of three photo-probes (diazirine-biotin, aryl-azide-biotin and phenol-biotin), each with a different labeling radius.^27^ Upon illumination with blue light, proteins proximal to the photocatalyst were biotinylated and captured by biotin enrichment using NeutrAvidin beads. Control samples with unconjugated lenacapavir and eosin Y showed that Gag-Cas9 (220 kDa), Cas9 (160 kDa), and mature capsid proteins were biotinylated and detected with all probes (Fig. 2e), because the eosin Y could diffuse throughout the particle. When eosin Y was conjugated to lenacapavir, preventing it from diffusing throughout the particle due to binding and localization at the capsid core, only the mature capsid protein, but not Cas9, was biotinylated regardless of the biotin probe used. The amount of the capsid protein that became biotinylated was proportional to the concentration of the lenacapavir-eosin Y conjugate used in the experiment. Each sample had similar quantities of input proteins, so the differences in abundance detected by biotin immunoprecipitation were caused by different localization of the photocatalyst and not sample loading. These results are consistent with our observations of uncleaved Gag-Cas9 in the EDVs^2,18^, where the Gag-Cas9 polyproteins are bound to the inner membrane of the EDVs and localized away from the capsid core. The prevalence of immature EDVs (Fig. 2c) also means that the majority of particles do not form a capsid core. Altogether, our data shows that it is unlikely that Cas9 associates with the core. The amount of Cas9 associated with the mature EDV capsid core was undetectable.

### EDV editing activity correlates with nuclear localization signal abundance on Cas9

Since the capsid core does not transport Cas9 RNPs to the cell nucleus, we reasoned that engineered NLSs on the Cas9 enzyme might be essential for nuclear entry and editing activity. To test this, we systematically removed both the heterologous N-terminal p53 and C-terminal SV40 NLSs from the packaged Cas9 enzymes (Fig. 3a). We observed a corresponding decrease in the luminescence of reporter cells treated with these EDVs, consistent with a requirement for NLS-mediated Cas9 nuclear transport. Removing the C-terminal SV40 NLS had a larger effect on reporter luminescence than removing the N-terminal p53 NLS. Removing all NLSs reduced the luminescence of EDV-treated reporter cells by more than 95%. We further tested whether the residual editing activity of the Cas9 RNPs without NLSs could be due to nuclear transport by the capsid core. Lenacapavir did not significantly decrease the luminescence of reporter cells incubated with EDVs packaging Cas9 RNPs lacking NLSs, indicating that the residual editing activity was not due to capsid core transport (Fig. 3b). These results show that the dominant route of nuclear delivery by EDVs is by NLSs rather than capsid core (Fig. 3c).

**Figure 3.**
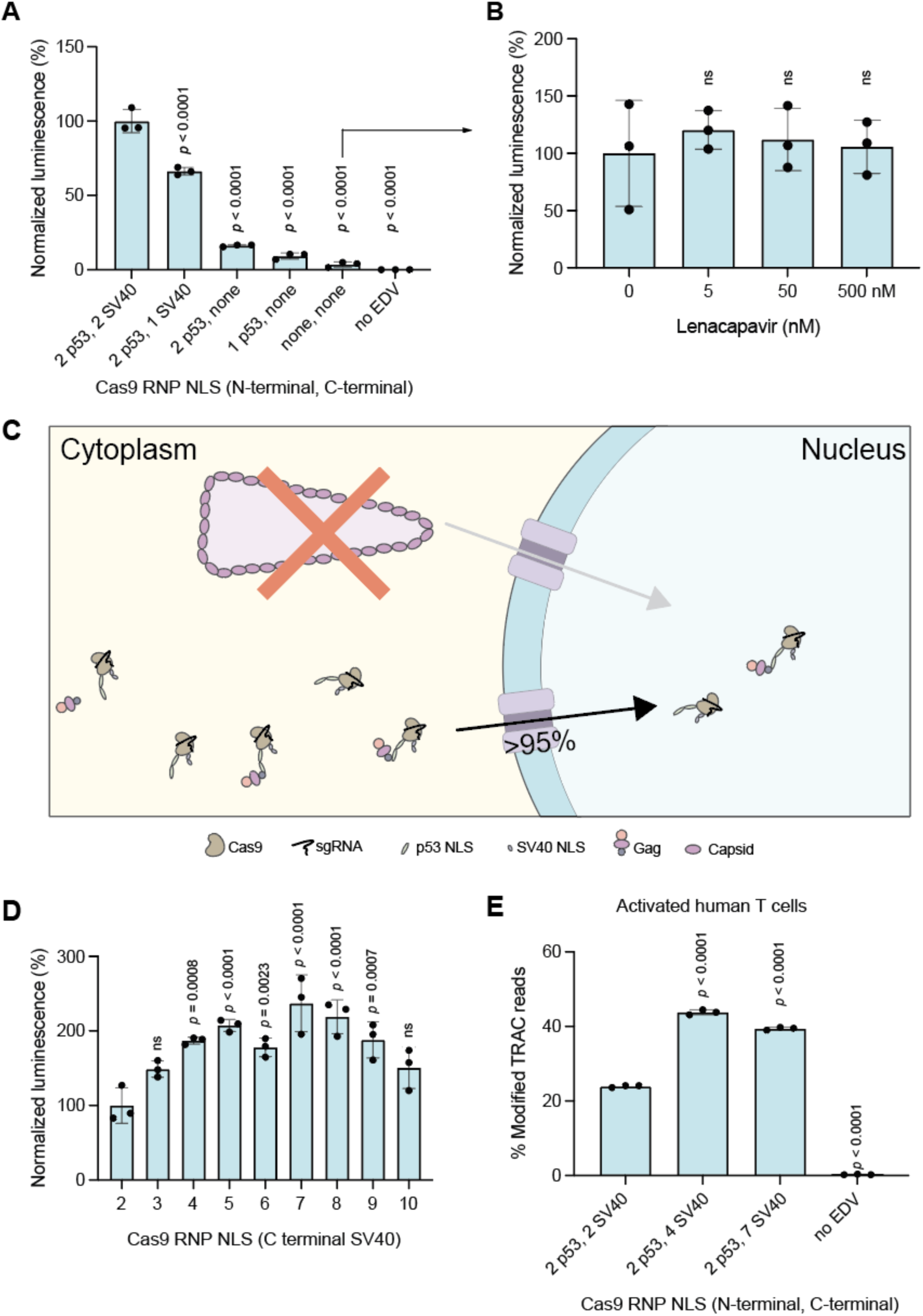
EDV editing activity correlates with nuclear localization signal abundance on Cas9. (a) Removal of NLS on Cas9 enzymes reduced editing and luminescence. Data were normalized to the current design with two N-terminal p53 and two C-terminal SV40 NLS. (b) The capsid core does not transport Cas9 enzymes missing nuclear localization signals (NLS) into the nucleus. EDVs packaging Cas9 enzymes with no NLS were incubated with luciferase reporter HEK-293T cells in the presence of Lenacapavir (0 - 500 nM). The luminescence of the reporter cells were recorded after incubation. (c) Summary schematic of Cas9 RNP nuclear delivery mechanism by EDVs. Cas9 RNPs are dominantly delivered to the nucleus by the NLS (bottom) and not by the capsid core (top). Schematic not to scale. Adding SV40 NLS to the C-terminus of RNPs in EDVs increased editing. (d) EDVs packaging Cas9 enzymes with two p53 N-terminal NLS and increasing numbers of SV40 NLS at the C-terminus were created. An equal volume of EDVs were incubated per design to capture both improvements in physical titer and per particle editing. Data were normalized to the current design (labeled as “2”). (e) EDV designs with four or seven NLS increased editing in activated primary human T cells as measured by amplicon sequencing at a dose of 4500 EDV per cell. The physical titers of the EDVs were determined using nanoparticle flow cytometry on a NanoFCM Nanoanalyzer instrument. *P*-values were calculated using an ordinary one-way ANOVA with Dunnett’s multiple comparisons test. Mean ± standard deviation of n = 3 biological replicates. Significant *p*-values as indicated. Non significance indicated by “n.s.”

We also found that adding additional NLSs to Cas9 enhances EDV-mediated Cas9 RNP editing efficiency. We created EDVs packaging Cas9 RNPs containing two to ten SV40 NLSs at the C-terminus of the Cas9 enzyme (Fig. 3d), because the C-terminal NLS had a larger effect on editing (Fig. 3a). Cas9s with four to nine NLSs showed ∼2-fold higher activity compared to the original two-NLS design, with seven NLSs being the best.^2^ Adding additional N-terminal NLSs to the Cas9 enzymes with seven C-terminal NLSs did not further improve EDV-mediated editing activity (*SI Appendix,* Fig. S4a). To confirm that the improvements in EDV editing were not specific to the luciferase reporter cells, we tested the four- and seven-NLS Cas9 designs in primary human T cells. We targeted the *TRAC* locus to disrupt the native T cell receptor (TCR), a step in the creation of therapeutic TCR-T cells.^28^ Activated T cells were incubated with an equal number of EDVs, and editing was quantified three days post-incubation by amplicon sequencing. EDVs packaging Cas9 RNPs with four or seven NLSs increased editing by 79% and 73% at the *TRAC* locus, respectively, compared to the original two-NLS designs (Fig. 3e). This increase in *TRAC* editing resulted in a corresponding reduction in the number of TCR-expressing T cells as quantified by flow cytometry (*SI Appendix,* Fig. S4b).

### Removing capsid core-related components created functional minimal EDVs

We wondered whether viral proteins that form or interact with the EDV capsid core could be removed, which could simplify particle production and avoid undesirable interactions with host cell proteins.^29,30^ Based on data showing that the C-terminal domain of the capsid protein was sufficient for Gag and Gag-pol proteins to assemble into immature HIV-1 virions,^22,31^ we removed the capsid N-terminal domain (amino acids 5 -148) or the entire capsid protein (amino acids 5 - 227) from the Gag and Gag-Pol polypeptides and tested the resulting EDVs in luciferase reporter cells. N-terminal domain removal had no effect but removing the entire capsid protein decreased editing by ∼75% (Fig. 4a). Next, we tested the removal of the Pol polyprotein, which includes viral protease, reverse transcriptase and integrase. The viral protease matures HIV-1 virions and may also liberate Cas9 RNPs from Gag proteins but is unnecessary in murine leukemia virus-based particles packaging base editors.^32^ Reverse transcriptase and integrase assemble with the capsid core to form the pre-integration complex for transgene integration. Removing either the viral protease (RT + INT) independently or the entire Pol polypeptide did not significantly decrease the activity of the EDVs, which shows that both Cas9 and Gag-Cas9 are functional (Fig. 4b). Similar deletion or truncation of the two remaining HIV-1 structural proteins, matrix and nucleocapsid, showed that matrix is required but nucleocapsid is not (Fig. 4c, d). Removing the nucleocapsid protein increased EDV-mediated editing by 43%. Altogether, these data show that most viral proteins related to the capsid core (N-terminal of the capsid, nucleocapsid, protease, integrase, and reverse transcriptase) are not necessary in EDVs.

**Figure 4.**
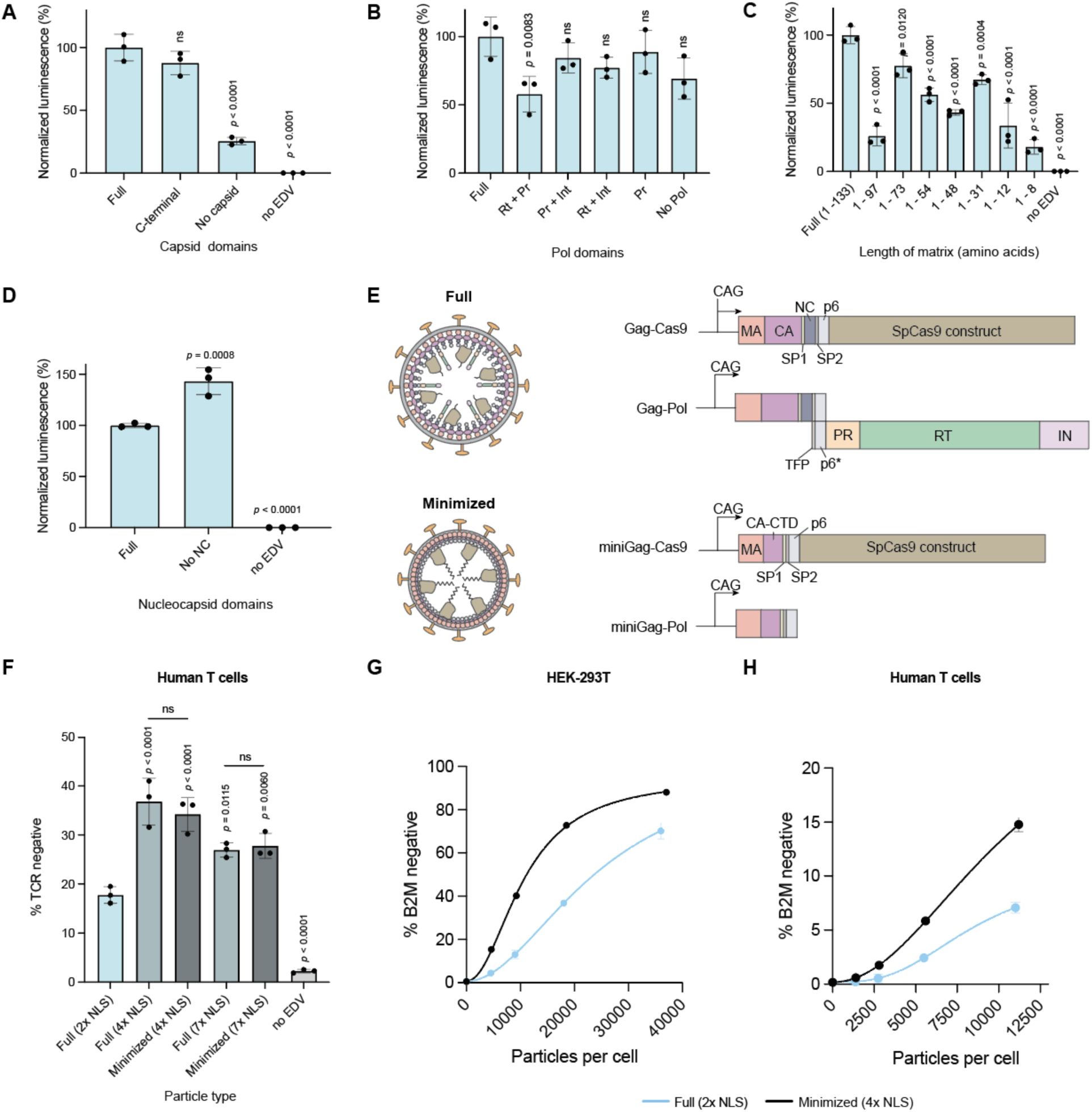
Removing capsid core-related components created functional minimal EDVs. (a) The N-terminal domain of the capsid protein (residues 5 - 149) or the entire capsid protein was removed and the activity of the EDVs was compared to EDVs containing the full capsid. An equal volume of EDVs were incubated per condition. Data were normalized to EDVs containing the full capsid. (b) Domains from the Pol polyprotein were removed systematically and the activity of the EDVs were compared to EDVs containing the full Pol polyprotein. Rt: Reverse transcriptase, Pr: Protease, Int: Integrase. An equal volume of EDVs were incubated per condition. Data were normalized to EDVs containing the full Pol. (c) The matrix protein was minimized one secondary structure element at a time from the C-terminal end. An equal volume of EDVs were incubated per condition. Data were normalized to EDVs containing the full matrix. (d) The nucleocapsid (NC) protein and the activity of the EDVs were compared to EDVs containing the full NC. Data were normalized to EDVs containing the full NC. (e) Schematic showing the components and plasmids involved in making full and minimized EDV. The minimized structural proteins are referred to as “miniGag” (f) Minimized EDVs had higher activity than the original full EDV design. Full, NLS optimized (7x and 4x) and minimized designs (7x and 4x) were incubated with primary activated human T cells (12000 particles per cell). Particle numbers were determined using the NanoFCM Nanoanalyzer. Expression of the T cell receptor was quantified five days after incubation. Mean ± standard deviation of n = 3 independent replicates. *P-*values were calculated using a one-way ANOVA with Šídák’s multiple comparison tests between the indicated pairs. Non-significance is indicated by “n.s.” (g) Editing at the *B2M* locus in HEK-293T cells was compared between minimized EDVs and full EDVs. Editing was determined five days after incubation using flow cytometry. Particle numbers were determined using the NanoFCM Nanoanalyzer. (h) Editing at the *B2M* locus in activated T cells was compared between minimized EDVs and full EDVs. Editing was determined five days after incubation using flow cytometry. Particle numbers were determined using the NanoFCM Nanoanalyzer. Unless stated otherwise, Mean ± standard deviation of n = 3 independent replicates. *P-*values were calculated using a one-way ANOVA with Dunnett’s multiple comparisons. Non-significance is indicated by “n.s.”

These deletions were combined to create minimal EDVs (miniEDVs) from both the four- and seven-NLS Cas9 designs due to their similar editing potency (Fig. 3). Although miniEDVs contain only 22% of the viral protein residues present in the original EDVs, they were produced with similar physical titers to the original particles (Fig. 4e; *SI Appendix,* Fig. S5a). Cryogenic electron tomograms showed these particles to be 25% smaller than the original EDVs (*SI Appendix,* Fig. S5b), with visible lipid envelopes and glycoproteins but lacking capsid cores (*SI Appendix,* Fig. S5c). Patches of protein density underneath the membrane may correspond to the minimized Gag protein (*SI Appendix,* Fig. S5d). The number of packaged Cas9 enzymes dropped from 391 ± 34 to 265 ± 16 per particle in original versus miniEDVs (*SI Appendix,* Fig. S5e), roughly consistent with the reduction in volume from ∼1.0×10^6^ nm^3^ to ∼4.3×10^5^ nm^3^ per particle. Notably the number of sgRNAs per particle remained consistent at ∼200 per particle (*SI Appendix,* Figure. S5f), suggesting that both original and miniEDVs package a similar number of Cas9 RNPs. We further found that miniEDVs could be produced with high functional titers without supplementing producer cells with plasmids encoding extra structural proteins (*SI Appendix,* Fig. S6), simplifying their production. In addition, single-chain antibodies can be displayed on their surface to mediate cell entry (*SI Appendix,* Fig. S7).

Lastly, we compared the editing efficiency of the miniEDVs to both our original and NLS-optimized EDV designs in primary human activated T cells at the *TRAC* locus by quantifying the decrease in T cell receptor expression by flow cytometry five days post-incubation (Fig. 4f). MiniEDVs packaging four- or seven-NLS Cas9s increased editing by 107% and 53%, respectively, relative to the original EDVs and were comparable to their respective NLS-optimized EDV counterparts. We further benchmarked the editing efficiency of the miniEDVs packaging four-NLS Cas9s across a range of concentrations in both HEK-293T cells and activated primary human T cells at the *B2M* locus, whose disruption enables production of allogeneic CAR T cells.^33^ We observed an average increase in editing per EDV particle of ∼2.5-fold in both HEK-293T and activated T cells. Thus, understanding the components inside of EDVs necessary for Cas9 delivery allowed us to simultaneously increase editing potency while streamlining the production of delivery vehicles for genome editing.

## Discussion

Virally-derived particles, including EDVs, have emerged as promising delivery vehicles for genome editing. We found that EDV-mediated Cas9 RNP editing activity is independent of the viral capsid core and instead depends on Cas9 nuclear localization signals (NLSs). Based on this finding and data from EDVs lacking additional viral proteins, we developed miniEDVs that showed 2.5-fold higher editing potency, relative to original EDVs, in both cell lines and primary cells. MiniEDVs can be produced in cells transformed with two plasmids, compared to three or more plasmids required for original EDVs.

Cryo-electron tomography showed that EDVs and lentiviral vectors share similar morphology and capsid structures. However, unlike lentivirus, EDVs do not use the internal capsid core for nuclear delivery of Cas9 RNPs. Instead, EDV-mediated genome editing depended on the presence of nuclear localization signal (NLS) peptides engineered onto Cas9. We also found that Cas9 RNPs are not associated with the capsid core, paving the way for removal of the capsid structure from EDVs to create simpler and more efficacious miniEDV particles.

The miniEDVs are 25% smaller than the original EDVs, yet packaged an equivalent quantity of guide RNAs and by extension Cas9 RNPs. The components of our minimized particles hold important lessons for engineering cell-derived particles and show that a membrane-binding domain (MA), an assembly domain (C-terminal CA), and a budding signal (p6) are sufficient for packaging and exporting a proteinaceous therapeutic cargo. Our miniEDVs do not contain functional viral enzymes (protease, reverse transcriptase or integrase) or viral nucleic acid-binding domains (nucleocapsid), reducing the possibility of unwanted interactions with target cells.

While we focused on an HIV-1-derived particle system, we anticipate that other virally-derived particles, including those based on related retroviruses, contain unnecessary proteins and could be simplified. As VLP and EDV systems advance toward clinical use, ensuring that these delivery vehicles contain only necessary components is critical to reduce complexity, improve manufacturing pipelines, and potentially reduce immunogenicity. The finding that miniEDVs require fewer plasmids to be produced while exhibiting higher editing activity and programmable cell entry, underscores the value of a mechanism-based approach to development. These results lay the groundwork for creating fully synthetic particles that use viral proteins to facilitate delivery, making genome editing therapies simpler, easier to produce and more efficacious.

## ACKNOWLEDGEMENTS

We thank all current and former members of the Doudna laboratory for their thoughtful input on this project, particularly Jennifer Hamilton, Connor Tsuchida, Kevin Wasko, Hannah Karp, Abdullah Syed, Karen Zhang, Nate Price, Katarzyna Soczek and Jason Nomburg. We thank D. Toso and R. Thakkar at the Cal-Cryo EM facility at QB3-Berkeley for help with EM data acquisition; P. Tobias and K. Stine for computing support. We would also like to acknowledge equipment and financial support from the James B. Pendleton Charitable trust

## AUTHOR CONTRIBUTIONS

Conceptualization: WN, JAD

Methodology: WN, JAD, JP, EN, RRS, ATS, ZL, JAW

Investigation: WN, JP, AB, ZWX, SH, RRS, ZL, ATS, RS, EN, JAD, JAW

Visualization: WN, JP, ZL, JAD

Funding acquisition: JAD, EN, RS, ATS, JAW

Project administration: WN, JAD

Writing and editing: WN, JP, AB, ZWX, SH, RRS, ZL, ATS, RS, EN, JAD, JAW

## DATA AND MATERIALS AVAILABILITY

Flow cytometry and imaging raw data files are available upon request. All other data are available in the main text or the supplementary materials. Plasmids generated in this study will be deposited on Addgene.

