## Supporting Information for "Mechanism-guided engineering of a minimal biological particle for genome editing"

**This PDF file includes:**

Materials and Methods
Figures S1 to S9
Tables S1 to S2
Legends for Movies S1 to S2
SI References

**Other supporting materials for this manuscript include the following:**

Movies S1 to S2

**MATERIALS**

HEK-293T cells were a gift from Melanie Ott. We purchased 15-cm plates (FB012925), 24-well plates (353047) and 10-cm plates (353003) from Fisher Scientific. DMEM (10-013-CV), fetal bovine serum (97068-085) and 0.45-µm filters (76479-020) were purchased from VWR. We purchased 1×PBS (21-040-CM), and trypsin (25-053-CI) from Corning. Penicillin-streptomycin (10,000 units/mL) (15140122), Opti-MEM™ I reduced serum medium ( 31985062), RIPA lysis and extraction buffer (89900), Halt™ protease inhibitor cocktail (100X) (78429), Pierce™ BCA protein assay kits (23225), SuperSignal™ west femto maximum sensitivity substrate (34094), Pierce™ ECL Western blotting substrate (32109), Restore™ PLUS Western blot stripping buffer (46428), NP-40 Surfact-Amps™ detergent solution (85124), Dynabeads™ human T-activator CD3/CD28 (11161D), custom TaqMan™ small RNA assays (CTRWFGP for luciferase sgRNA, CTZTEYN for *CLTA* sgRNA, CTCE4RX for *TRAC* sgRNA, and CTWCW3V for the *B2M* sgRNA), MicroAmp™ optical 384-well reaction plates (4309849), DAPI (4',6-diamidino-2-phenylindole, dilactate) (D3571), NeutrAvidin agarose beads (29200), One Shot™ Mach1™ T1 phage-resistant chemically competent *E. coli* (C862003) and NuPAGE™ LDS sample buffer (4X) (NP0007) were purchased from Thermo Fisher Scientific, Inc. We purchased 2-mercaptoethanol (M6250), DL-dithiothreitol (DTT) (D9779), anti-HIV-1 p24 antibody produced in rabbit (SAB3500946), monoclonal anti-Flag M2 antibody produced in mouse (F1804), ethylenediaminetetraacetic acid disodium salt dihydrate (ED2SS), magnesium chloride (M4880), bovine serum albumin (A2058), copper (II) sulfate (C1297), sodium L-ascorbate (A4034), and dimethyl sulfoxide (DMSO) (D8418) from Sigma-Aldrich. DNA/RNA shield DirectDetect™, ZymoPURE II plasmid midiprep (D4200), ZymoPURE II plasmid maxiprep (D4202), ZymoPURE II plasmid gigaprep (D4204) and Mix & go *E. coli* transformation kit & buffer sets (T3002) were purchased from Zymo Research Corporation. Sucrose (S24060) was purchased from Research Products International. Primers and oligonucleotide sequences were purchased from Integrated DNA Technologies. Plasmids pJRH-1179 U6-reci Gag-Cas9 v2 (Add gene 201915) and pJRH-1180 U6-reci Gag-pol v2

(Addgene 201914) were a gift from Jennifer Hamilton. Plasmids pMD2.G (12259), psPAX2 (12260), pHAGE-CMV-Luc2-IRES-ZsGreen-W (164432), and U6-sgRNA-EFS-Cas9-P2A-Puro (211687) were ordered from Addgene. NEBuilder® HiFi DNA assembly master mix (E2621X), Luna® universal one-step RT-qPCR kit (E3005E), nuclease-free water (B1500L), thermolabile proteinase K (P8111S) and Q5® high-fidelity 2X master mix (M0492L) were purchased from New England Biolabs. EM gold tracer, 10 nm (25487), and Quantifoil R2/2, UT, 200 mesh, gold grids (Q2100AR2-2nm) were purchased from Electron Microscopy Sciences. Polyethylemimine, Linear (MW 25,000) (23966) was purchased from Polysciences. HIV1 p24 ELISA kits (ab218268) were purchased from Abcam. Criterion TGX stain-free gel 4-20% (5678094), precision plus protein kaleidoscope standards (1610375), mini bio-spin columns (7326207), and nitrocellulose membranes, 0.2 µm (1620112) were purchased from Bio-Rad Laboratories, Inc. IRDye® 800CW goat anti-rabbit IgG secondary antibody (926-68070), IRDye® 800RD goat anti-mouse IgG secondary antibody (926-32210) and IRDye® 680RD goat anti-mouse IgG secondary antibody (926-32211) were purchased from Li-COR Biosciences. Goat anti-mouse IgG heavy and light chain cross-adsorbed antibody HRP conjugated (A90-516P) and goat anti-rabbit IgG heavy and light chain cross-adsorbed antibody HRP conjugated (A120-201P) were purchased from Fortis Life Sciences. Mouse anti-cas9 antibody (61758) was purchased from Active Motif, Inc. EZH2 (D2C9) XP® rabbit mAb #5246 (5246S) was purchased from Cell Signaling technology. GAPDH antibody (G-9) (sc-365062) was purchased from Santa Cruz Biotechnology. Lenacapavir (HY-111964), and PF74 (HY-120072) were purchased from MedChemExpress. Passive lysis buffer (E1941), and CellTiter-Glo 2.0 cell viability assays (G9242) were purchased from Promega Corporation. CulturPlate-96 black plates (6005660) were purchased from PerkinElmer. Illumination™ lyophilized firefly luciferase enhanced assay kit (I-935-1000) was purchased from Gold Biotechnology. Human peripheral blood leukopak (200-0092), and EasySep™ human T cell isolation kits (100-069) were purchased from STEMCELL Technologies. X-VIVO 15 serum-free hematopoietic cell medium (04-418Q) was purchased from Lonza. N-acetyl-L-cysteine USP

(VWRV0108-25G) was purchased from VWR International, LLC. Recombinant human IL-7 (200-07), and recombinant human IL-2 (200-02) were purchased from PeproTech, Inc. Silica quality control nanospheres were purchased from NanoFCM, Inc. Recombinant human IL-15 (247-ILB) was purchased from R&D systems. Cas9 ELISA kits (PRB-5079) were purchased from Cell Biolabs. APC anti-human β2-microglobulin antibody (316312) and APC anti-human TCR α/β antibody (306718) were purchased from BioLegend, Inc. We purchased a 10% SDS solution (S0288) from Alpha Teknova, Inc. QIAprep spin miniprep kits (27106), QIAquick gel extraction kits (28704) and QIAquick PCR purification kits (28104) were purchased from Qiagen. SPRIselect beads (B23318) were purchased from Beckman Coulter Life Sciences. PhiX sequencing control v3 (FC-110-3001), NextSeq 500/550 mid output kit v2.5 (150 Cycles) (20024904) and HT1 hybridization buffer (20015892) was purchased from Illumina Inc. Isothiocyanate-EY (90091) was purchased from Biotium. Azide-PEG$_4$-amine (BP-21615) was purchased from BroadPharm. BTTAA (1236-100) was purchased from Click Chemistry Tools (Vector Laboratories).

**METHODS**

*Cell culture.* HEK-293T cells (1.6 million) were seeded into 15-cm plates in complete DMEM media (DMEM with 10 v/v% FBS and 1 v/v% penicillin/streptomycin, 20 mL). Cells were incubated at 37°C and 5% $CO_2$ for 72 h and passaged as required. HEK-293T cells were routinely checked for mycoplasma infection (Stem Cell Core, Gladstone Institutes). T cells were isolated from human peripheral blood Leukopaks using the EasySep™ isolation kit following the manufacturer's instructions. Deidentified donors (< 60 years old, non-smoking) were chosen without regard to sex, gender, ethnicity and race. T cells were used fresh for *in vitro* experiments without freezing. For activation, T cells were seeded (1,000,000/mL) in complete X-VIVO 15 media (5 v/v% FBS, 4 nM N-acetyl-cysteine, and 55 μM 2-mercaptoethanol) and activated with anti-human CD3/CD28 magnetic Dynabeads (1:1 beads to cells) for 24 h with IL-2 (200 units/mL) in complete X-VIVO 15 media. After activation the magnetic beads were removed and the T cells were cultured in X-VIVO 15 medium with FBS (5 v/v%),  IL-7 (10 ng/mL), IL-15 (5 ng/mL), and IL-2 (200 units/mL).

*Plasmids.* CRISPR-Cas9 spacer sequences are shown in Supplementary Table 1. The appropriate spacer sequences were cloned into U6-sgRNA-EFS-Cas9-P2A-Puro, pJRH-1179 U6-reci Gag-Cas9 v2 (referred to as Gag-Cas9) and pJRH-1180 U6-reci Gag-pol v2 (referred to as Gag-Pol) plasmids using NEBuilder® HiFi DNA assembly. For constructing the luciferase reporter cell lines, a C205ATC mutation was generated in pHAGE-CMV-Luc2-IRES-ZsGreen-W by ordering the appropriate primers and HiFi DNA assembly.  Deletions (nuclear localization signals or Gag-Pol domains) were made to the Gag-Cas9 and Gag-Pol plasmids using NEBuilder® HiFi DNA assembly. Nuclear localization signals were inserted into the Gag-Cas9 plasmids using NEBuilder® HiFi DNA assembly of the appropriate plasmid fragments, where the sequences of the nuclear localization signal with appropriate overhangs were purchased as double-stranded oligonucleotides from IDT.  Plasmids were transformed into Mach1 *E.coli* cells rendered competent using the Mix & Go *E. coli* transformation kit following the manufacturer's instructions. Mach1 cells were grown with the appropriate antibiotic selection. Plasmids were extracted and purified using mini-, midi-, maxi-, or giga- prep kits as necessary following the manufacturer's instructions. All plasmids were sequence-verified by whole plasmid sequencing (Primordium Labs and Plasmidsaurus Inc.) before use.

*Cryogenic electron tomography (cryo-ET) of EDVs and lentiviral vectors.* HEK-293T cells (4 million) were seeded into 10-cm plates and allowed to attach overnight. For Enveloped Delivery Vehicle (EDV) production, the Gag-Pol (3300 ng), Gag-Cas9 (6700 ng), and VSV-G mut (1000 ng) plasmids were diluted in Opti-MEM and mixed with polyethylenimine (PEI) (3:1 PEI: plasmid ratio by mass). The mixture was incubated for 30 min at room temperature and added dropwise

(~400 µL total) to the HEK-293T cells. Cells were incubated with the transfection mixture for at least 6 h, then swapped with Opti-MEM (10 mL). EDVs designed to edit luciferase were used for safety, as the spacer does not have complementary sequences in the human genome. The mutant VSV-G defective for binding was used for similar reasons. Lentiviral vectors were produced similarly using the Gag-Pol (10000 ng) and VSV-G mut (1000 ng) plasmids. No transgene encoding plasmid was included for safety. Particles were collected the following morning (~48 h after transfection). Cell debris was removed by centrifugation (4000 *g* for 10 min) and filtering through a 0.45-µm filter. Particles (~30 mL) were concentrated by iodixanol cushion (10 w/v% OptiPrep in 1× PBS) ultracentrifugation (100,000g for 75 min). The supernatant was removed and the pellets were resuspended in 1× PBS (0.1 mL). Samples were kept on ice and used freshed. EM grids (QF AU200 R 2/2, Quantifoil) were plasma-treated using a Tergeo-EM plasma cleaner (Pie Scientific). The purified particles (2 µL) were mixed with 10-nm gold fiducials (1 µL) and applied to EM grids at 4°C and 90% humidity inside a Vitrobot Mark IV (Thermo Fisher Scientific) and allowed to adsorb for 30 sec. Samples were blotted (3 - 5 sec, blot force 5) then plunge frozen in liquid ethane. Collection of cryo-ET data was performed as previously published.[1] Tilt series were collected on a Titan Krios G3i 300kV cryogenic transmission electron microscope (Thermo Fisher Scientific) with a K3 direct electron detector and an energy filter (Gatan) with a pixel size of 1.67 Å. Tilt series were acquired from -60° to 60° in 3° increments using a dose-symmetric tilt scheme, a defocus range of -2 to -4.5 µm, and a total dose of 120 electrons/Å$^2$. At each tilt position the total exposure was split into 4 frames. Detailed imaging parameters per dataset are summarized in Supplementary Table 2. Tilt images were motion-corrected (Motioncorr2) and exposure filtered based on the accumulated dose using the Alignframes function in IMOD 4.11.[2] The contrast transfer function (CTF) for each tilt image was determined on non-exposure filtered tilt images using CTFFIND4.[3] Tilt series were subsequently aligned using the gold fiducials using Etomo in IMOD 4.11. For visualization, tomograms were reconstructed by weighted back projection and isotropically binned by a factor of four, then three dimensionally gaussian filtered by 1 - 2.5 standard deviations in Matlab (Mathworks). Tomograms were visualized in either IMOD 4.11 or UCSF Chimera 1.18. We manually annotated the particles as immature (concentric densities underneath the bilayer), mature (internal core structure), and unknown. Slices through tomograms were exported using IMOD 4.11 and cropped using Adobe Illustrator 2024. The diameters of the particles were measured in IMOD 4.11 across the vertical and horizontal central axes and averaged to determine the particle size. Graphs were plotted in GraphPad Prism v.10.1.1 (270).

*Subtomogram averaging.* For subtomogram averaging, unbinned tilt series were 3D-CTF corrected based on the determined defocus values and subsequently reconstructed by weighted projection using NovaCTF.[4] Particle centers and radii were manually determined for immature particles using the volume tracer function and the Pick Particle plug in UCSF Chimera 1.18 or UCSF Chimera 1.15.[5] Subtomogram averaging for EDVs and lentiviral vectors was performed as previously described using a combination of Matlab scripts (MATLAB R2023B, Mathworks) based on functions from the TOM, AV3 and Dynamo packages and subTOM 1.1.6 (https://github.com/DustinMorado/subTOM).[1,6–8] Initial subtomogram positions and orientations were determined based on the radius and the particle center and were sampled along the surface of a sphere at on the level of the particle membrane. Overlapping subtomograms with an edge length of 428 Å were extracted from the 3D-CTF corrected tomograms binned 4× times. An initial reference was generated by iteratively aligning subtomograms from a single tomogram using an exhaustive search. This process resulted in a low resolution average that resembled previously determined structures of the immature HIV capsid protein displaying 6-fold symmetry. The reference and corresponding subtomograms were shifted to center the 6-fold symmetry axis. Subtomograms were then re-extracted from the updated positions and 6-fold symmetry was applied during subsequent iterations of subtomogram alignment until the reference did not improve further. This average was then supplied as a reference to align the full data set from the

4× binned tomograms. The full data set was split into two half sets by particle and both half sets were aligned independently from each other with identical parameters. The full data set was aligned for 5 iterations with 6-fold symmetry applied and subtomogram positions converged onto overlapping positions. Subtomograms with low cross correlation values were subsequently removed. Once the reference and resolution did not improve further, subtomograms with an edge length of 390 Å were re-extracted at the aligned positions from 2× binned tomograms. After 6 iterations of alignment, the resolution and quality of the map did not improve further. The resolution was determined by calculating the Fourier Shell Correlation (FSC) between the two independently aligned half sets. Existing models of HIV-1 immature capsid (PDB: 5L93) were fitted by rigid body fitting into the final, sharpened and filtered map using the fit-in-map functionality in UCSF chimera 1.15. For lattice map analysis and visualization, the 3D positions and orientations of each subtomogram (corresponding to each aligned CA hexamer) were displayed back into the original tomogram using the "Place Object" plug-in[9] in UCSF Chimera 1.15 and quantified per EDV or lentiviral vector in Matlab.

*Western blot analysis of EDVs and lentiviral vectors.* For comparing the amount of Gag to capsid, EDVs and lentiviral vectors were produced as above. EDVs (1 mL) were harvested 30, 48 and 72 h after transfection and frozen at -80°C. The concentration of capsid domains in each sample was determined using a capsid (p24) ELISA kit and normalized across samples using RIPA buffer. Samples (30 μL) were mixed with 4× LDS (10 μL with 5 v/v% 2-mercaptoethanol) and loaded into the wells of a Criterion TGX Stain-free gel. We ran the SDS-PAGE gel (100 V, 1 h, room temperature), then transferred proteins in the gel to nitrocellulose membranes (40 V, overnight, 4°C). Membranes were blocked (1 h, room temperature) in a blocking buffer (5% Non-fat dry milk and 0.1 v/v% Tween-20 in 1× tris-buffered saline). Rabbit anti-p24 antibody and mouse anti-Flag antibody (1/2500 dilution in blocking buffer) were added and incubated (overnight, 4°C). Blots were washed three times with 1× TBST (0.1 v/v% Tween-20 in 1× tris-buffered saline). IRDye®

680RD goat anti-mouse IgG secondary antibody and IRDye® 800CW goat anti-rabbit IgG secondary antibody (1/500 dilution in blocking buffer) was added and incubated (1 h, room temperature). Blots were imaged using a ChemiDoc™ MP Imaging System (Bio-Rad Laboratories, Inc.) after washing three times with 1× TBST. Images were inverted and processed in ImageJ2 V. 2.14.0 and cropped in Adobe Illustrator 2024 for presentation. For determining the nuclear association of the Cas9 enzymes and the capsid proteins, EDVs were made as described above and incubated (7 mL) with HEK-293T cells in 15-cm dishes with PF74 at the indicated concentrations. DMSO was used as a vehicle control. After 24 h, the cells were trypsinized, washed with 1×PBS, then re-suspended in 1×PBS (1 mL). Some of the cells (100 µL) were taken, pelleted by centrifugation (100 *g*, 2 min) and lysed in RIPA with protease inhibitors (30 min, on ice). The samples were centrifuged (20,000 *g*, 10 min, 4°C) and the supernatant was transferred to a new tube and stored as the "total" fraction. The remaining cells (900 µL) were pelleted by centrifugation and lysed in cytoplasm lysis buffer (315 µL, 10 mM Tris-HCl pH 7.4 with 1 mM DTT, 1 mM $MgCl_2$, 10% sucrose, 100 mM NaCl, 0.5 v/v% NP-40 and 1× protease inhibitors) on ice. After 10 min, the samples were centrifuged (664 *g*, 2 min, 4°C) and the supernatant was transferred to a new tube and stored as the "cytosolic" fraction. The pellet was washed by centrifugation (20,000 *g*, 10 min, 4°C) using cytoplasm lysis buffer (1 mL) three times, then resuspended in RIPA with protease inhibitors. The nuclei were lysed (30 min, on ice), and insoluble components were removed by centrifugation (5,200 *g*, 2 min, 4°C). The supernatant was transferred to a new tube and stored as the "nuclear" fraction. The protein concentration of the samples were quantified using BCA assays and normalized to 1 µg/µL, then mixed with the appropriate amount of 5 v/v% 2-mercaptoethanol in 4× LDS. Samples were heated (5 min, 90°C) then loaded (30 µL) into a 4-12% PAGE gel. SDS-PAGE gels were run, then transferred and blocked as described above. The blots were first stained with either mouse anti-cas9 antibodies (1/100 in blocking buffer) or mouse 1/100 anti-Flag antibodies for 48 h at 4°C. Blots were washed three times with 1× TBST then incubated with HRP-conjugated goat anti-mouse secondary antibodies (1/5000 in blocking buffer) for 1 h at room temperature. Blots were washed three times with 1× TBST. SuperSignal™ West femto maximum sensitivity substrate was subsequently added and the blots were imaged using a ChemiDoc™ MP Imaging System (Bio-Rad Laboratories, Inc.). After imaging, blots were stripped using a stripping buffer following the manufacturer's manual, then blocked as described above. Blots were then stained for capsid protein using a rabbit anti-p24 antibody (1/1000) for 2 h at room temperature, washed three times with 1× TBST, the stained with HRP-conjugated goat anti-rabbit secondary antibodies (1/5000 in blocking buffer). Blots were washed and imaged as described above using Pierce™ ECL Western Blotting Substrate. The procedure was repeated to stain for EZH2 (1/2000 rabbit anti-EZH2 in blocking buffer) and GAPDH (1/5000 mouse anti-GAPDH in blocking buffer). Band intensities were quantified using the measure function in ImageJ2 V. 2.14.0, then normalized to the band intensity of DMSO conditions. Images were processed as above for presentation. The raw images of the blots are shown in Supplementary Figure 9.

*Proximity ligation experiments.* EDVs (30 mL) were produced and concentrated by ultracentrifugation using a sucrose cushion (20 w/v%, 2 mL). Prior to bioconjugation, we examined the structure of lenacapavir bound to capsid and found that the alkyne group was not necessary for drug binding.[10] Lenacapavir-eosin Y (EY) was made using a two-step bioconjugation procedure. Isothiocyanate-eosin Y (SCN-EY, 1.0 equiv.) was coupled with azide-PEG$_4$-amine (N$_3$-amine, 1.1 equiv.) in freshly made sodium bicarbonate (10 mM) for 2 h at room temperature in the dark to generate azide-eosin Y (azide-EY). A mixture of BTTAA (100 µM), copper sulfate (80 µM) and lenacapavir (1.2 equiv., 400 µM) was mixed in sodium ascorbate acid (200 µM) and then immediately added to the azide-EY mixture. The reaction was incubated at 37°C for 3 h to generate lenacapavir-EY. EDVs were incubated (room temperature, 15 min) with lenacapavir-EY at a range of concentration (50 - 500 nM). For the control experiments, EDVs were incubated with unconjugated lenacapavir and unconjugated eosin Y (500 nM each). The appropriate photoprobes (diazirine-biotin, aryl-azide-biotin or phenol-biotin) were added to the EDVs (100 μM final concentration) and incubated at room temperature for 5 min. The samples were then illuminated (100% intensity, 10 min) using a Penn PhD Photoreactor M2 (Sigma Aldrich, Z744035) with a 450 nm blue light source module (Sigma Aldrich, Z744033). Fan speed was set at 6800 rpm under manual control with 100 r/min stirring. The EDVs were then lysed in 1× RIPA buffer with protease inhibitors on ice for 30 min. The samples were centrifuged (20,000 g, 10 min, 4°C) and the supernatant was transferred to a new tube with 10% of the sample saved as "input" samples. LDS sample buffer (1× final) was added to the "input" samples. Biotin immunoprecipitation was performed on the remainder of the samples. In brief, NeutrAvidin agarose beads (50 μL) were washed three times (0.5 mL, 1× PBS) then added to the EDV lysates and incubated (4°C, 16 h). The samples were loaded on mini bio-spin columns and the flowthrough was discarded. The beads were washed three times with 1× RIPA lysis buffer (0.5 mL), three times with NaCl solution (1M NaCl in 1× PBS, 0.5 mL), then three times with urea solution (2M urea freshly dissolved in 50 mM ammonium bicarbonate, 0.5 mL). The beads were subsequently suspended in LDS sample buffer (2×, 40 μL) and boiled (95°C, 5 min). For immunoblotting, proteins were run on 4-12% gels, then transferred from SDS-PAGE gels to PVDF membranes using an iBlot-2 dry blotting system (Thermo Scientific, IB21001). Membranes washed with Tris buffered saline (TBST, 37mM sodium chloride, 20mM Tris, 2.7mM potassium chloride, 0.05% Tween 20; pH=7.4) and blocked with BSA solution (TBST containing 0.1% Tween-20 and 5% BSA) before the antibodies (1/1000) were added as indicated. Immunoblot images were captured by an infrared LI-COR imager (LI-COR Biosciences, Odyssey CLx). Data were analyzed and visualized using ImageStudioLite (v5.2.5) and Adobe Illustrator (v22.1).

*Luciferase assays.* Luciferase HEK-293T cells were made by transducing low passage HEK-293T cells with a lentiviral vector packaging a mutant luciferase (C205ATC) and zsGreen transgene. Monoclonal cell lines were established by sorting the cells for zsGreen expression using a BD

FACSAria Fusion cell sorter (BD Biosciences). For small molecule drug experiments, lenacapavir and PF74 were dissolved in DMSO. EDVs editing the luciferase transgene were produced as described above. Lentivirus were made using the Gag-Pol (10000 ng), pMD2.G (1000 ng) and transgene plasmid U6-sgRNA-EFS-Cas9-P2A-Puro with the luciferase gRNA (2500 ng). Luciferase HEK-293T cells (6000 cells per well) were incubated with lentivirus or EDVs (100 μL for PF74, 75 μL for lenacapavir) with the indicated concentrations of PF74 or lenacapavir in black bottom 96-well plates. An equivalent volume of DMSO was used as the vehicle control. After 48 h, the media from the luciferase HEK-293T cells incubated with EDVs was removed and replaced with a passive lysis buffer (1× in ultrapure water, 20 μL) and incubated on a rocking shaker (room temperature, 30 min). After incubation, the luminescence of the wells were recorded on a Tecan Spark multimode microplate reader (Tecan Group Ltd.) by injecting luciferase substrate (30 μL) into each well immediately before measurement. The conditions with lentiviral vectors were quantified as above 72 h after transduction to provide sufficient time for the transgene to integrate, produce Cas9 and edit the reporter. For experiments using EDVs without nuclear localization sequences, EDVs were produced with the appropriate Gag-Cas9 plasmid as described above. We transduced luciferase HEK-293T cells with EDVs (100 μL). The luminescence of the wells were measured 48 h after transduction as described above. To test whether EDVs without NLSs could use the capsid core for transport, EDVs packaging Cas9 RNPs without NLS were produced. EDVs (50 μL) were incubated with luciferase HEK-293T cells with the indicated concentrations of lenacapavir. The luminescence of the wells were recorded 48 h after transduction. For experiments using EDVs with more NLSs, the appropriate EDVs were produced and incubated (6.25 μL) with the luciferase HEK-293T cells. The luminescence was recorded 48 h after transduction. For experiments using EDVs with deletions in the Gag-Pol polypeptide, the appropriate EDVs were produced and incubated (capsid deletions 40 μL, Pol deletions 25 μL, matrix deletions 40 μL, nucleocapsid deletions 40 μL, combining deletions 10 μL) with luciferase HEK-293T cells. The luminescence was recorded 48 h after transduction. In all experiments, the luminescence signal was normalized by the vehicle control conditions or positive control conditions and plotted in GraphPad Prism v.10.1.1. Statistics as indicated were calculated in GraphPad Prism v.10.1.1.

*Characterization of EDVs.* The physical titers of the EDVs were quantified using a NanoAnalyzer instrument (NanoFCM) following the manufacturer's SOP. Briefly, EDVs were diluted in Tris-HCl buffer (100 mM Tris-HCl pH 7.5 with 1 mM EDTA) at least ten-fold before analysis. Silica quality controls beads of known concentration were used as calibration standards to determine the physical titre. Particle concentrations were determined using instrument software, NanoFCM Profession V2.0. For quantifying the sgRNA or Cas9 quantity inside the particles, the particles were first purified by ultracentrifugation, then diluted in either DirectDetect buffer for reverse transcription quantitative polymerase chain reactions (RT-qPCR) or RIPA buffer for ELISA. For RT-qPCR, Custom TaqMan™ small RNA assays were designed and ordered against the gRNA sequence. Synthetic sgRNA sequences with the appropriate spacers and no modifications were used as standards for quantification. Samples (4 µL) or standards (4 µL) were mixed with RT-qPCR mix (6 µL) (1× Luna Luna® Universal One-Step RT-qPCR mix with 0.25× RT primer and 1× small RNA assay probes from TaqMan™ small RNA assay kit in nuclease-free water). The entire sample or standard was loaded into 384-well plates. Samples and standards were prepared in duplicate. RT-qPCR was performed on a QuantStudio™ 5 Real-Time PCR System (Thermo Fisher Scientific, Inc.) with the following parameters: carryover prevention (25°C, 30 sec), reverse transcription (55°C, 15 min), initial denaturation (95°C, 1 min), and 45 cycles of denaturation (95°C, 10 sec), extension (60°C, 60 sec) with a plate read. Data was processed in Microsoft Excel Version 16.83. For Cas9 or p24 enzyme-linked immunosorbent assays, samples and standards were diluted in RIPA as appropriate. The manufacturer's instructions were followed. Data was processed in Microsoft Excel Version 16.83. All data was plotted and statistics calculated as indicated in GraphPad Prism v.10.1.1.

*Editing in HEK-293T cells.* The appropriate EDVs editing *B2M* were produced as described above. The physical titers of the particles were determined using the NanoAnalyzer instrument as above. HEK-293T cells (2000 cells per well) were incubated with EDVs at the indicated concentrations in 24-well plates and incubated at 37°C. The cells were trypsinized 5 d after transduction and washed with a staining buffer (1× PBS with 2 w/v% bovine serum albumin). The cells were stained (on ice in the dark, 30 min) with APC anti-human β2-microglobulin antibody (5 µL per sample) and DAPI as a viability stain (0.3 nmol per sample) in the staining buffer. The cells were then washed three times with the staining buffer. Flow cytometry was used to quantify B2M expression using an Attune Nxt Flow Cytometer (Thermo Fisher Scientific, Inc.). Data was analyzed in FlowJo 10.9.0 with the gates shown in Supplementary figure 8. All data was plotted and statistics calculated as indicated in GraphPad Prism v.10.1.1.

*Editing in primary activated T cells.* Activated T cells (30,000 for NLS experiments, 10,000 for EDV minimization experiments) were incubated with EDVs (concentrations as indicated). For flow cytometry, T cells were harvested 5 d after incubation, and stained for B2M as above. We stained for expression of the T cell receptor similarly, but using an APC anti-human TCR α/β antibody (5 µL per sample). Flow cytometry was used to quantify expression using an Attune Nxt Flow Cytometer (Thermo Fisher Scientific, Inc.). Data was analyzed in FlowJo 10.9.0 with the gates shown in Supplementary figure 4. For next generation sequencing, T cells were harvested 3 d after incubation, then spun down (300 *g*, 3 min). The cell media was removed and the cells were lysed in lysis buffer (10 mM Tris-HCl pH 7.5 with 0.05 w/v% SDS and 0.72 U/mL of thermolabile proteinase K, 200 µL). The proteinase K was subsequently deactivated by incubation at 55°C for 20 min. Extracted DNA was stored at -80°C until use. The *TRAC* loci was amplified from gDNA (1 ng) by TRAC-ngs.fwd and TRAC-ngs.rev primers (0.5 µM each) (Supplementary Table 1) using NEB Q5 PCR mastermix (1×). PCR was performed using the Applied Biosystems ProFlex PCR system (Thermo Fisher Scientific, Inc.) with the following parameters: initial denaturation (95°C, 3 min), 25 cycles of denaturation (95°C, 10 sec), annealing (65°C, 20 sec) and extension (72°C, 30 sec), and a final extension (72°C, 60 sec). This first PCR product was purified using SPRIselect beads (0.8×) following the manufacturer's protocol. A second PCR was performed to ligate illumina P5 and P7 adapter and indexing sequences for sequencing (Supplementary Table 1). The product of the first PCR (1 ng) was mixed with forward and reverse primers (0.5 μM each) and NEB Q5 PCR mastermix (1×). PCR was performed using the Applied Biosystems ProFlex PCR system (Thermo Fisher Scientific, Inc.) with the following parameters: initial denaturation (95°C, 3 min), 10 cycles of denaturation (95°C, 15 sec), annealing (60°C, 20 sec) and extension (72°C, 30 sec), and a final extension (72°C, 60 sec). The product of the second PCR was gel purified using the QIAquick gel extraction kit following the manufacturer's protocol. PCR products were checked for purity and the correct size using the 2100 Bioanalyzer (Agilent Technologies, Inc.). The second PCR products were pooled to create a library (1 nM of each product). PhiX was added (30 v/v% of 1 nM). The library was denatured with sodium hydroxide (0.1N final concentration) for 5 min at room temperature, then TRIS-HCl (pH 7, 67 mM final concentration) was added. The library was diluted (1.5 pM final concentration) in HT1 Hybridization Buffer before loading onto a NextSeq 500/550 Mid Output Kit v2.5. Sequencing was performed on a Illumina NextSeq 500 sequencer (Illumina, Inc.). To quantify editing efficiency from our NGS experiments we used CRISPResso2 using default parameters for an NHEJ run as previously published (https://github.com/pinellolab/CRISPResso).[11] For batch characterization we followed standard parameters for quantification of editing efficiency by CRISPResso2 as outlined at the github link above. Editing rates were reported as the maximum indel frequency as determined by CRISPResso2.

**SUPPLEMENTARY FIGURES**

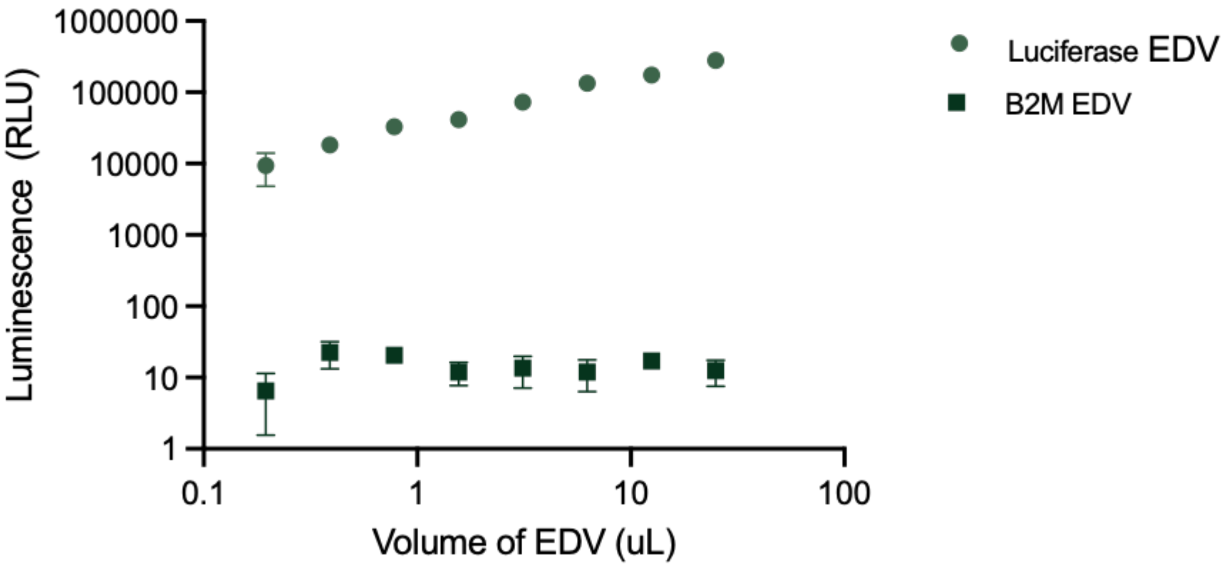


**Supplementary Figure 1.** Luminescence from the reporter cell line was linear with the volume of EDVs incubated (circles). EDVs that do not edit the truncated luciferase locus do not turn on the reporter cell line (squares). Mean ± range of n = 2 biological replicates.

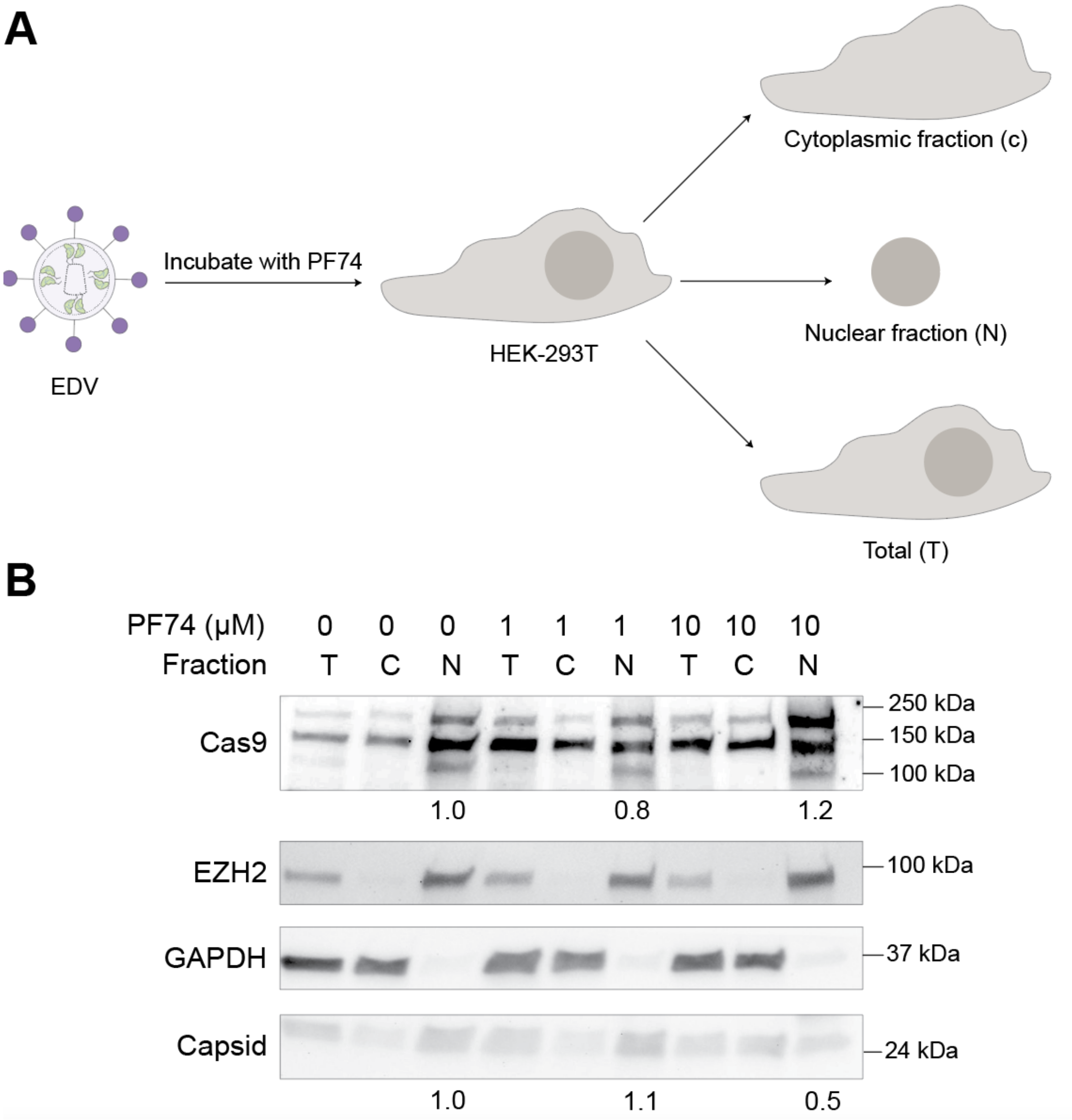


**Supplementary Figure 2.** Western blot showing inhibition of capsid core nuclear entry does not reduce Cas9 entry into the nucleus. (a) HEK-293T cells were incubated with EDVs in the presence of PF74 inhibitors (0 - 10 µM). Cells were collected and fractionated into total, cytosolic and nuclear fractions 24 h of incubation. (b) Western blot of the fractions. EZH2 and GAPDH were used as nuclear and cytosolic markers respectively. The intensity of the Cas9 and capsid protein were measured in ImageJ, normalized to the DMSO control (0 nM), and indicated in the relevant lanes. The experiment was repeated twice with similar results.

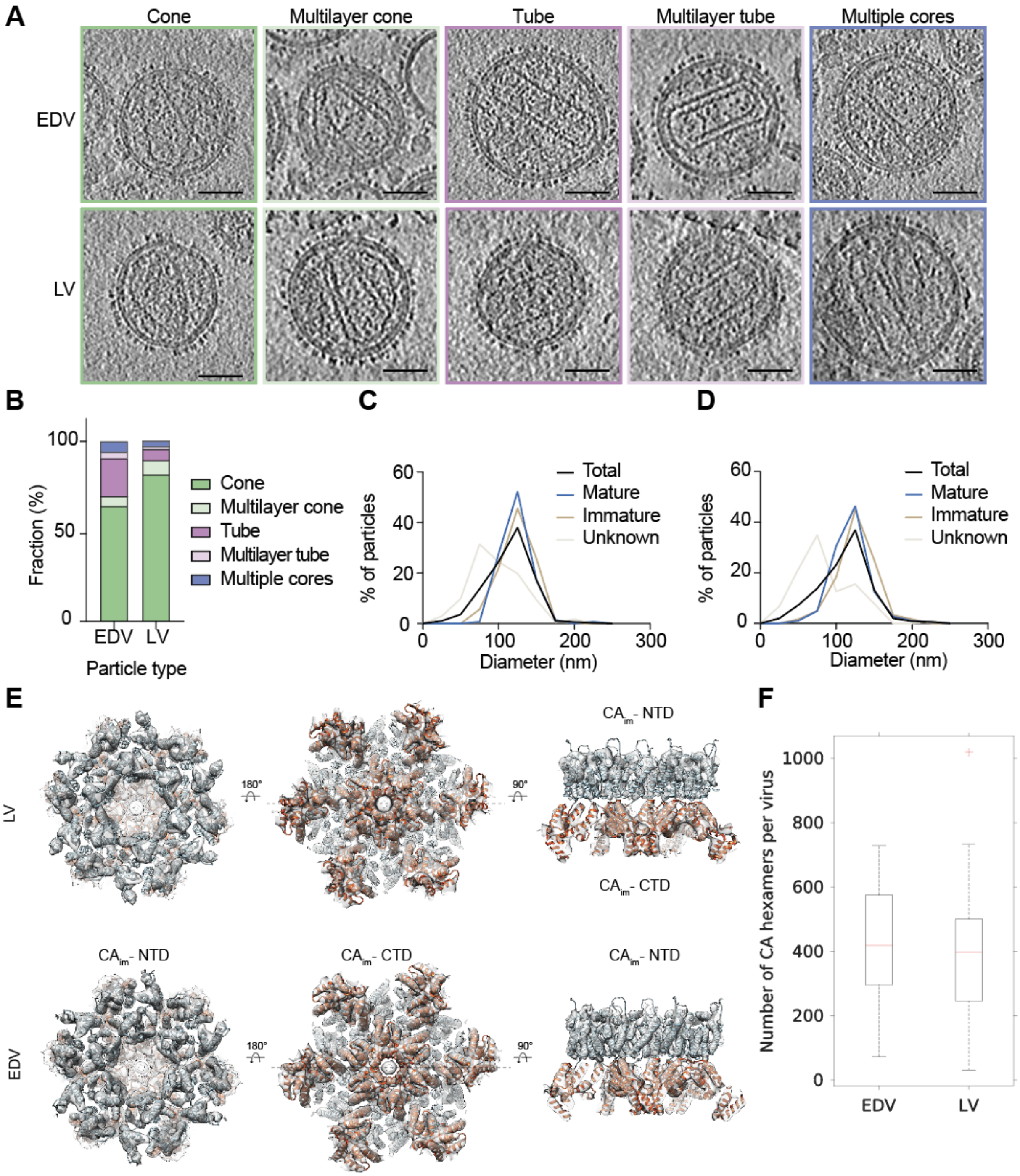


**Supplementary Figure 3.** Structural characterization of EDVs and lentiviral vectors (LVs) using cryo-ET. (a) Representative two-dimensional slices in the XY plane of the cryo-tomograms of EDVs and LVs containing different core morphologies. Scale bars are 50 nm. (b) Fraction of mature EDVs (n = 142) and LVs (n = 189) that have conical, tubular, multilayer or multiple cores. (c) Size distribution of EDVs (measured by the diameter of the outer lipid membrane) N = 498. (d) Size distribution of lentivirus. N = 374. (e) Cryo-ET 3D reconstructions of immature capsid from LVs and EDVs obtained by subtomogram averaging in gray at 10 Å resolution and 9 Å resolution, respectively. HIV-1 CA structures were fitted into the gray density maps (PDB 5L93). The density shown on the left is oriented with the CA- N-terminal domain (NTD) facing upward, the density map in the middle with CA- C-terminal domain (CTD) facing upwards. On the right, a cross section through the density at the position indicated by the dashed line in the middle panel is shown. The NTD regions of the fitted CA models are colored in blue and CTD regions of the fitted CA models are colored in orange. (f) Number of CA hexamers per EDV or lentiviral particle. The cross indicates an outlying datapoint.

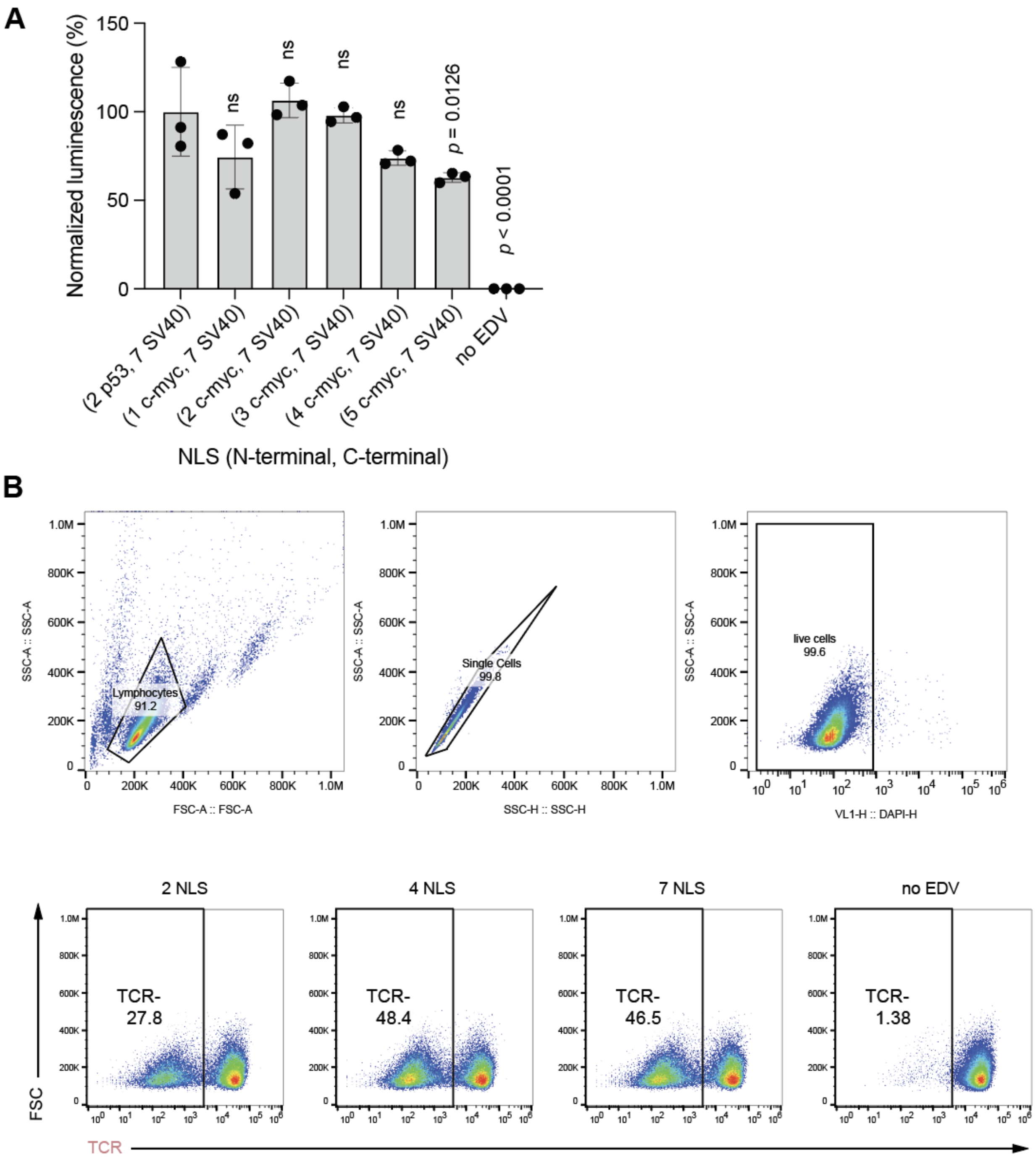


**Supplementary Figure 4.** Characterization of EDVs with more NLS. (a) Adding additional N-terminal NLS did not improve EDV editing. To facilitate tiling of NLS, we first replaced the bipartite p53 NLS (18 amino acids) with a monopartite c-myc NLS (9 amino acids). Both NLS use the same mechanism of nuclear import.[12,13] EDVs were then incubated with the luciferase HEK-293T cells. (b) Flow cytometry quantification shows that four and seven NLS Cas9 RNPs in EDVs decreased the number of primary activated human T cells expressing the TRAC protein in concordance with the sequencing results. The gating strategy is shown in the top panels. One representative biological replicate of three is shown.

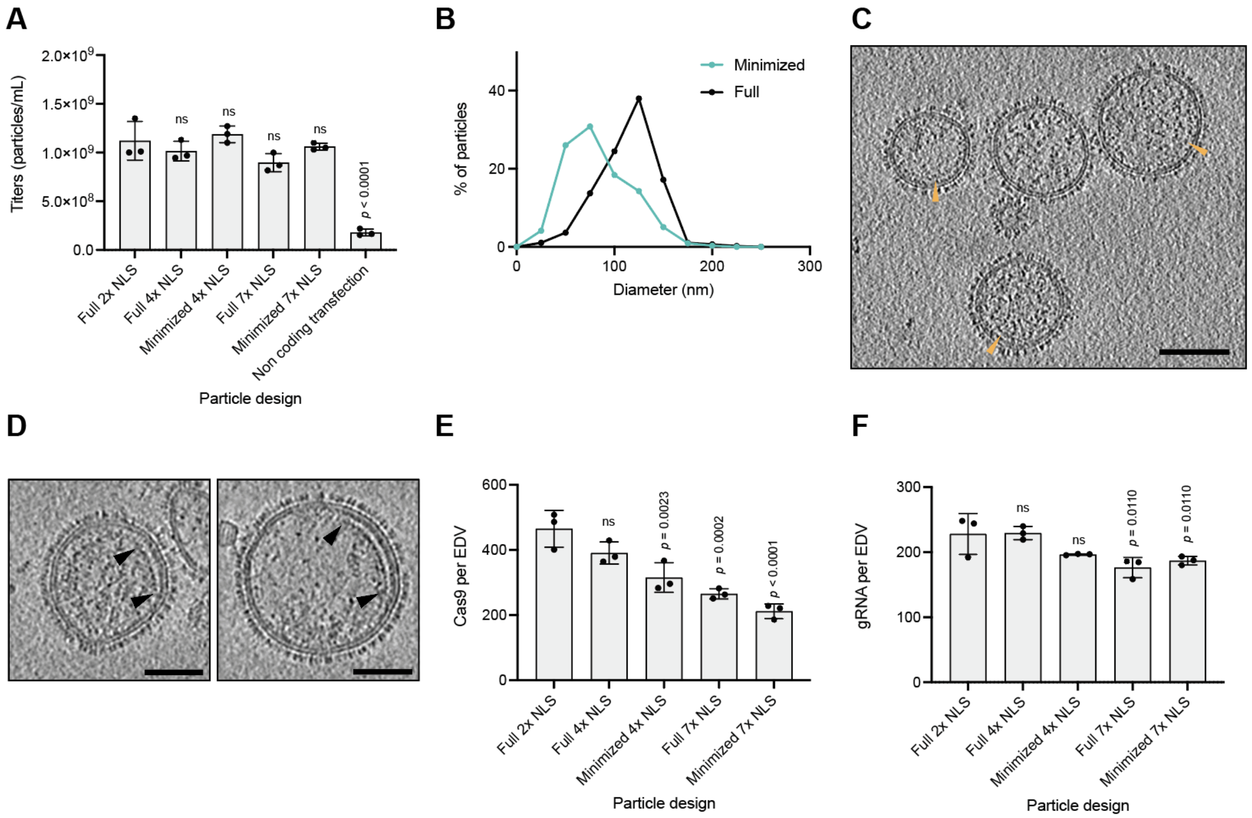


**Supplementary Figure 5.** Characterization of miniEDVs. (a) Quantification of the physical titers of EDVs using the NanoFCM NanoAnalyzer. (b) Size distribution of miniEDVs (N = 315) compared to full EDVs (N = 498) measured as the diameter of the outer lipid membrane in cryo-tomograms. (c) Two-dimensional slice of a minimized EDVs tomogram. Scale bar is 50 nm. Orange arrows point to densities underneath the lipid bilayer that potentially corresponds to the minimized Gag. (d) Averaged 66.8 Å thick slice of minimized EDV to show the protein density underneath the lipid membrane (black arrow). Scale bar is 50 nm. (e) Quantitation of Cas9 enzymes per particle for EDVs targeting *TRAC* measured by ELISAs. The particle number was determined by nanoparticle flow cytometry. (f) Quantitation of gRNA per particle for EDVs targeting *TRAC* measured by RT-qPCR. The particle number was determined by nanoparticle flow cytometry.

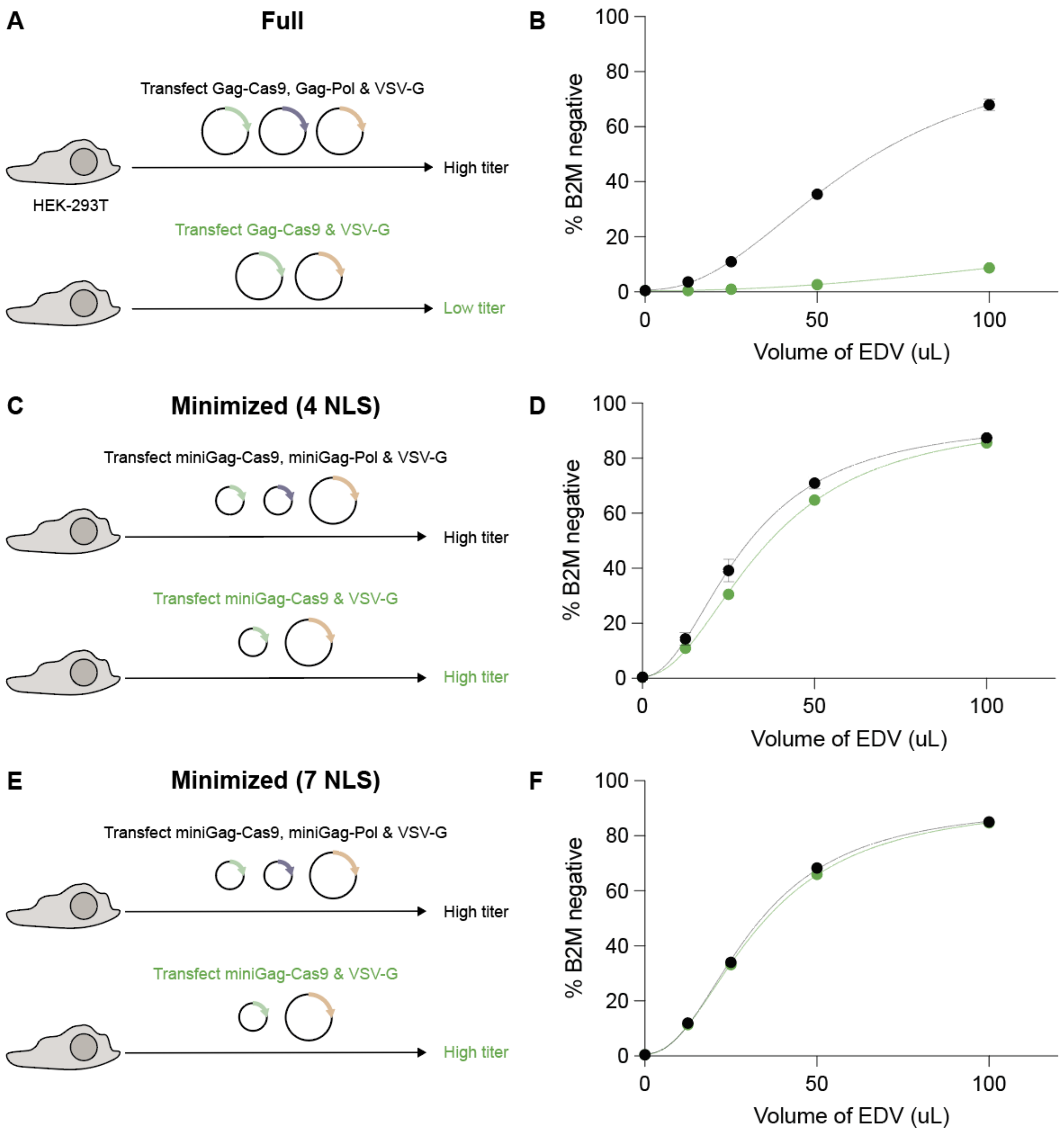


**Supplementary Figure 6.** MiniEDVs could be produced without supplementing producer cells with structural plasmids encoding extra Gag-Pol structural proteins. (a) Schematic illustrating the production of the full particles, which is efficient with a 2:1 mass ratio of Gag-Cas9 to Gag-Pol plasmids. (b) Producing EDVs using only the Gag-Cas9 plasmids results in low functional titers. Full EDVs were produced with either a 2:1 ratio of Gag-Cas9:Gag-Pol plasmids (black) or only Gag-Cas9 (green). EDVs were incubated with HEK-293T cells and the number of edited cells was quantified by flow cytometry 5 d after incubation. (c) Schematic illustrating the production of the miniEDVs with four NLS, which is efficient without the added miniGag-Pol plasmids. (d) MiniEDVs could be produced using only the miniGag-Cas9 plasmids with only a small decrease in functional titer. MiniEDVs were produced with either a 2:1 ratio of Gag-Cas9:Gag-Pol plasmids (black) or only Gag-Cas9 (green). (e) Schematic illustrating the production of the miniEDVs with seven NLS, which is efficient without the added miniGag-Pol plasmids. (d) MiniEDVs could be produced using only the miniGag-Cas9 plasmids with only a small decrease in functional titer. MiniEDVs were produced with either a 2:1 ratio of Gag-Cas9:Gag-Pol plasmids (black) or only Gag-Cas9 (green). EDVs were incubated with HEK-293T cells and the number of edited cells was quantified by flow cytometry 5 d after incubation. Mean ± standard deviation of n = 3 independent replicates. Data points were fitted to a sigmoidal curve for visualization.

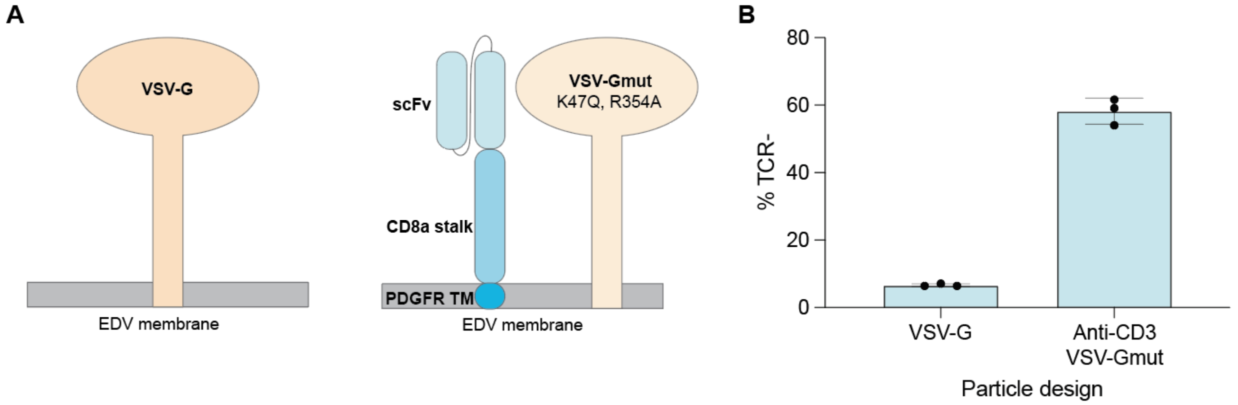


**Supplementary Figure 7.** Single-chain variable fragments (scFvs) can be displayed on the surface of miniEDVs to mediate cell entry. (a) Schematic of a vesicular stomatitis virus glycoprotein (VSV-G) pseudotyped EDV membrane and a EDV membrane displaying a scFV using a CD8a stalk and platelet-derived growth factor receptor (PDGFR) transmembrane (TM) domain. The EDV membranes displaying scFvs are supplemented with a receptor binding deficient mutant of VSV-G (VSVG-mut) that retains fusogen activity. (b) Editing activity of a scFv miniEDV and a VSV-G miniEDV in primary activated human T cells. T cell receptor (TCR) expression was quantified with flow cytometer five d after incubation. Mean ± standard deviation of n = 3 independent replicates.

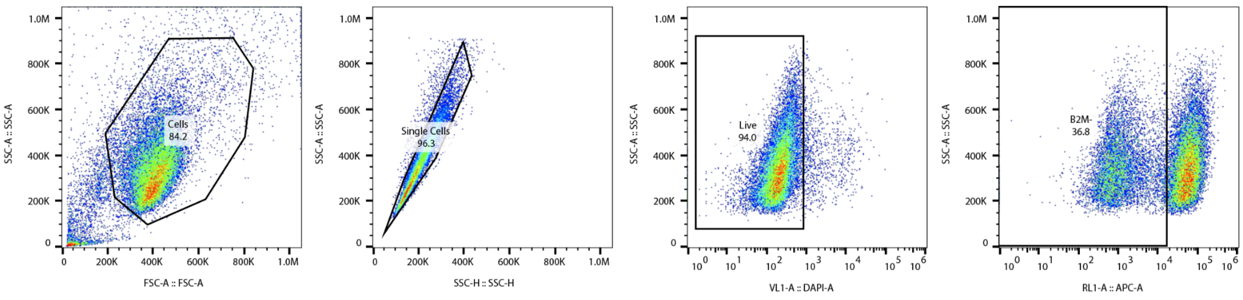


**Supplementary Figure 8.** The gating strategy for quantifying the expression of B2M in HEK-293T cells.

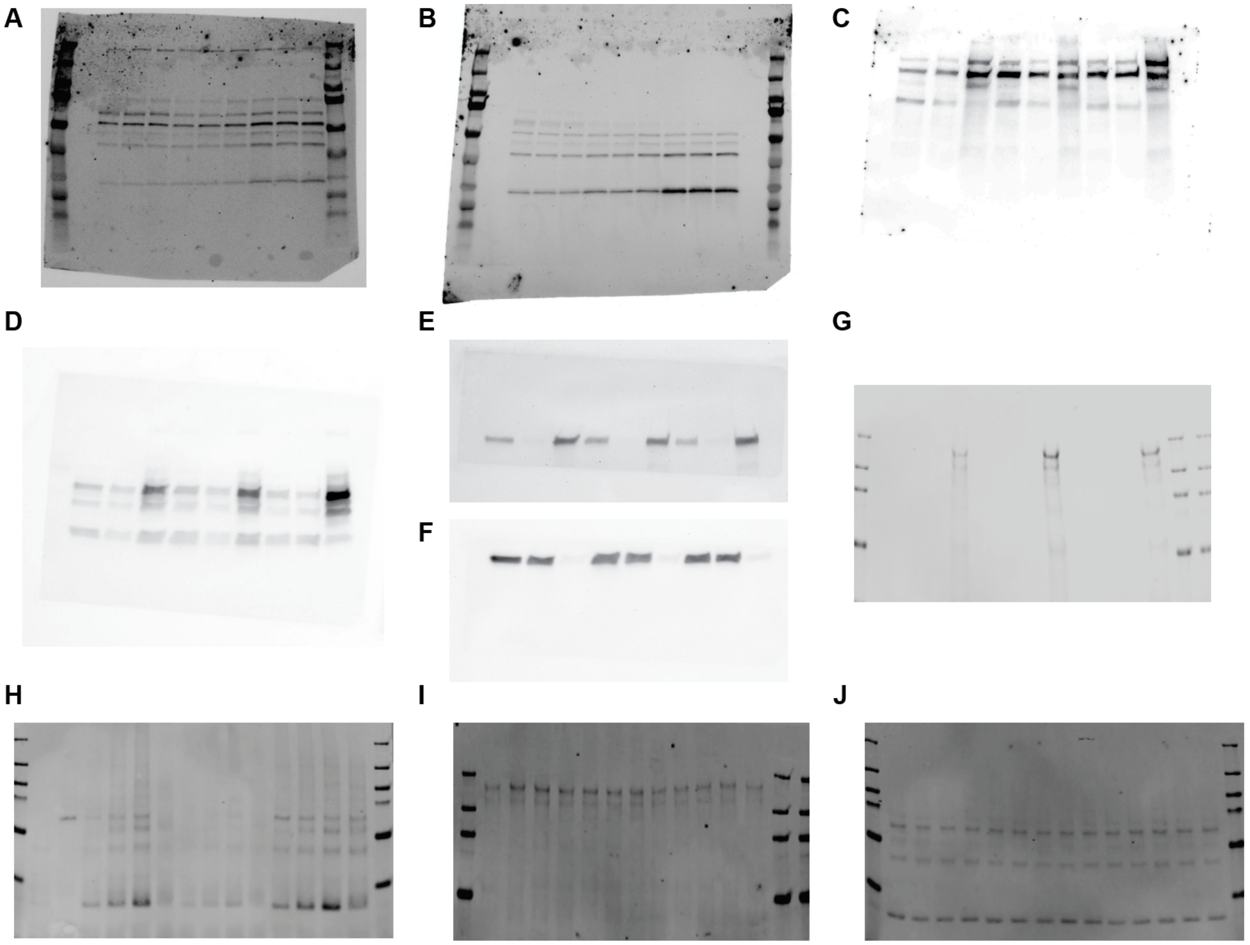


**Supplementary Figure 9.** Raw Western blots. (a) Blot from Figure 1c. EDVs were stained for the capsid domain. (b) Blot from Figure 1c. Lentiviral vectors were stained for the capsid domain. (c) Blot from Supplementary Figure 3. The cell fractions were stained for Cas9. (d) Blot from Supplementary Figure 3. The cell fractions were stained for the capsid domain. (e) Blot from Supplementary Figure 3. The cell fractions were stained for EZH2. (f) Blot from Supplementary Figure 3. The cell fractions were stained for GAPDH. (g) Blot from Figure 2e. The biotinylated proteins were stained for Cas9 using anti-Flag antibodies. (h) Blot from Figure 2e. The biotinylated proteins were stained for the capsid protein. (i) Blot from Figure 2e. The input proteins were stained for Cas9 using anti-Flag antibodies. (j) Blot from Figure 2e. The input proteins were stained for the capsid protein.

### SUPPLEMENTARY TABLES

**Supplementary Table 1: Oligonucleotide sequences**

| Name | Usage | Sequence (5' to 3') |
|---|---|---|
| Mutant Luciferase | CRISPR spacer | AGAAGCTATGAAAATCGATA (PAM: TGG) |
| TRAC | CRISPR spacer | CAGGGTTCTGGATATCTGT (PAM: GGG) |
| B2M | CRISPR spacer | GAGTAGCGCGAGCACAGCTA (PAM: AGG) |
| TRAC-ngs.fwd | Primers for NGS | ACACTCTTTCCCTACACGACGCTCTTCCGATCTGAGGGAAATGAGATCATGTCC |
| TRAC-ngs.rev | Primers for NGS | GTGACTGGAGTTCAGACGTGTGCTCTTCCGATCTATGTCTAGCACAGTTTTGTC |
| Illumina_P5 | Primers for NGS | AATGATACGGCGACCACCGAGATCTACAC**NNNNNNNN**ACACTCTTTCCCTACACGACGCTCTTCCGATCT |
| Illumina_P7 | Primers for NGS | CAAGCAGAAGACGGCATACGAGAT**NNNNNNNN**GTGACTGGAGTTCAGACGTGTGCTCTTCCGATCT |

Illumina adaptor sequences used for library prep are in green and index sequences are **bolded.**

**Supplementary Table 2: Cryo-EM data collection and processing parameters**

| | Immature CA EDV | Immature CA LV | Streamlined EDVs |
|---|---|---|---|
| **Data collection and processing** | | | |
| **Magnification** | 53,000 X | 53,000 X | 53,000 X |
| **Voltage (kV)** | 300 | 300 | 300 |
| **Detector** | K3 | K3 | K3 |
| **Energy-filter** | Yes | Yes | Yes |
| **Slit width (eV)** | 20 | 20 | 20 |
| **Pixel size (Å)** | 1.68 | 1.68 | 1.68 |
| **Defocus range (µm)** | -2 to -4.5 | -2 to -4.5 | -2 to -4.5 |
| **Defocus step (µm)** | 0.25 | 0.25 | 0.25 |
| **Tilt range (min/max, step)** | -60/60°, 3° | -60/60°, 3° | -60/60°, 3° |
| **Tilt scheme** | Dose-symmetrical | Dose-symmetrical | Dose-symmetrical |
| **Total Dose (electrons/Å²)** | 120 | 120 | 120 |
| **Frame number** | 4 | 4 | 4 |
| **Tomograms used for structure/acquired** | 38/47 | 28/49 | |
| **Final # subtomograms (set A/ set B)** | 20064/22302 | 8161/8024 | |
| **Symmetry imposed** | C6 | C6 | |
| **Final assymetric units** | 254196 | 97110 | |
| **B-factor** | -1200 | -1500 | |
| **Global resolution (Å)** | 9.3 | 10.3 | |

### SUPPLEMENTARY MOVIES

**Supplementary Movie 1.** Representative tomogram of EDVs.

**Supplementary Movie 2.** Representative tomogram of lentiviral vectors.
